# Intuitive knowledge of object acoustics enables perceptual separation of physical variables from impact sounds

**DOI:** 10.64898/2026.01.28.702236

**Authors:** Vinayak Agarwal, James Traer, Jeremy Schwartz, Josh H. McDermott

**Author notes:** co-first authors (Vinayak Agarwal, James Traer).

## Abstract

Upon hearing objects collide, humans can estimate physical attributes such as material and mass. Although the physics of sound generation is well established, the inverse problem that listeners solve – of inferring physical parameters from sound – remains poorly understood. Classical accounts posit the use of acoustic cues that correlate with physical variables, but do not explain how humans might distinguish multiple concurrent physical causes. To study this problem, we built a probabilistic generative model of impact sounds, combining theoretical acoustics with statistics of object resonances measured from hundreds of everyday objects, and used it to synthesize and manipulate experimental stimuli. Humans accurately judged object properties from collision sounds. However, when both of the colliding objects varied, performance was impaired if the distribution of object resonances deviated from those measured in real-world objects. The results suggest that listeners use internal physical models to separate the acoustic contributions of objects in the world.

## Introduction

When two objects collide, they create a sound, from which humans can readily judge their physical properties. Consider for example, a plate set on a table, or a glass marble bouncing on a wood board. Everyday experience suggests that a listener can identify classes of objects (e.g. a plate, a ball), motions (e.g. impacting and settling, bouncing and rolling) and materials (e.g. ceramic, wood, glass)^1^. Each of these judgments can be construed as the solution to an inverse physics problem, as the sound of an impact results from the structure and motion of the colliding objects via physical laws.

The creation of sound by objects is a classic problem in physics^2–4^. Objects are known to vibrate most strongly at certain frequencies known as “resonant modes”^5,6^, the frequencies, powers and decay rates of which are determined by the object shape and material^7^. In recent years, high-resolution acoustical models have allowed the synthesis of realistic impact sounds for a wide range of complex objects and motions^8–11^, provided that the object geometry and material can be specified with sufficient precision.

By contrast, the inference of physical properties from sound remains a challenging problem^12^, and its solution by the human brain remains poorly understood. Humans are able to estimate some physical parameters from impact sounds, such as material^13–20^, size^21–24^, and force of impact^25^, and can in some cases discern aspects of an object’s shape^22,26–28^. However, the means by which humans make such judgments remains unclear.

The most commonly articulated explanation of auditory physical estimation abilities is that humans have learned to associate particular acoustic cues with physical variables. For instance, the hardness of a struck object has often been linked to the ratio of the decay rate of sound to its frequency^14,20,22^, or to a combination of decay rate, frequency and amplitude^29^. Similarly, the mass of a dropped object has been linked to spectral centroid and overall sound level^23,24,27^.

One challenge for cue-based explanations is that sound is always produced by the interaction of multiple physical variables, typically involving multiple objects. For instance, consider an object dropped on a surface (Fig. 1a). The sound that results is a function of the resonant properties of the surface^5,6^, but also of the force created upon impact^25,30^, which depends on the height from which the object was dropped, its mass^23^, and the stiffnesses of the object and surface. As a result, it is not obvious that any single cue can be diagnostic of any single physical variable, because the cue is likely to be influenced by the other concurrent physical variables.

**Figure 1.**
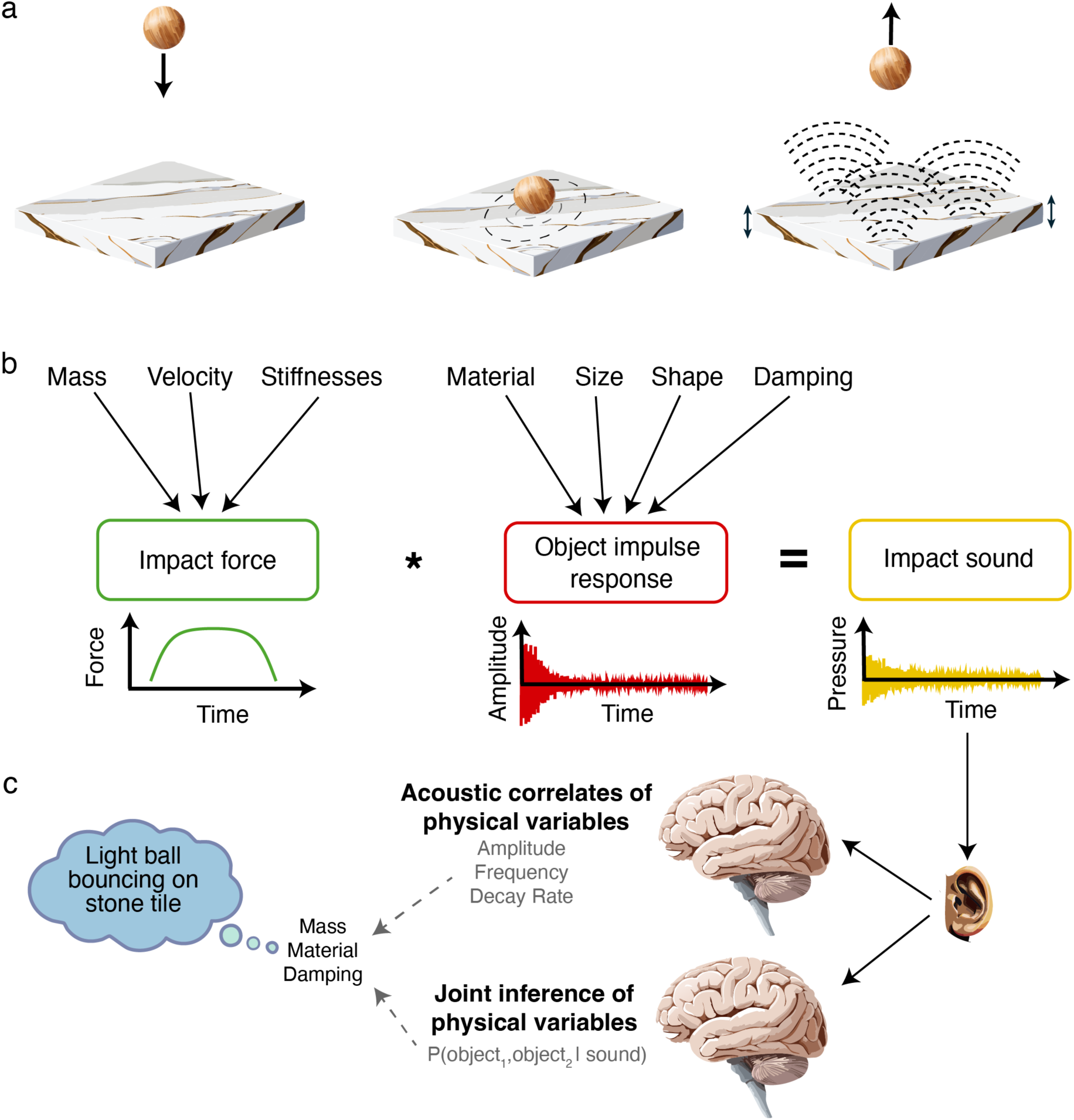
Physics of impact sounds. **a**. Impact sound produced by a ball striking a surface. When the ball contacts the surface, a momentary force is created as the surface and ball compress and then separate. This force induces vibrations in the surface that resonate within it and then radiate into the surrounding medium, causing the sound that we hear. Vibrations are also induced in the ball, but if the ball is small these may produce negligible sound in comparison to the struck surface. **b**. Linear systems model of impact sound generation, with dependencies between physical variables and sound generation. An impact sound can be approximated as the convolution of a time-varying impact force and the impulse response of the vibrating surface. The force is a function of the ball mass and velocity as well as the stiffnesses of the ball and surface. The impulse response of the surface is a function of its properties, including material, size, shape, and damping. The presence of these multiple causal influences on the resulting sound renders the problem of inferring the properties of either object ill-posed. **c**. Alternative explanations of human physical inferences. A common proposal (top) is that humans have learned acoustic correlates of physical variables, and use these to estimate these variables. An alternative (bottom) is that humans jointly infer (i.e., perceptually “separate”) the properties of the underlying objects, using some type of knowledge of how sound is generated from physical variables.

An alternative perspective on physical estimation is that estimating any individual physical factor requires “separating” its causal contribution from that of the other physical factors that determine a sound. Under this view, physical inferences present the auditory system with an ill-posed scene analysis problem analogous to other problems that necessitate the separation of multiple causal variables, such as the “cocktail party problem”^31,32^ in which a listener must separate a single voice from a mixture of voices, or to the problem of separating environmental reverberation from the sound of a voice in a room^33^. Thus far the inference challenges imposed by concurrent physical variables have remained poorly understood and rarely studied. Much previous research on the perception of physical variables from sound has considered settings in which only one physical variable is varied within an experiment, leaving it unclear a) whether humans can judge physical variables in realistic settings in which multiple physical causes of sound must be disambiguated, and b) how humans might overcome the ill-posed nature of the underlying inference problem.

Ill-posed inference problems in perception are typically solved using implicit knowledge of the regularities of the underlying causal factors, as these regularities narrow the space of possible solutions. We hypothesized that humans have internalized physics-induced regularities in how sound is generated, and use this intuitive knowledge to jointly infer the multiple physical causal factors that produce any single sound. Analogous hypotheses are commonly discussed in other domains of perception^34^, but have not been widely considered in the domain of intuitive physics. To test this hypothesis, we first sought to characterize the physical structure underlying impact sounds.

Although classical physics provides a general framework for this understanding^2,3,5,6^, it is not sufficiently specified to make reliably accurate predictions about the everyday objects that humans typically encounter (see Discussion). We thus began by conducting a large-scale acoustic characterization of over 400 everyday objects. We measured the resonances of each object, revealing systematic acoustic effects of material, size, mass, and damping. We incorporated these empirically measured statistical effects into a physics-based generative model of impact sounds, allowing realistic sounds to be synthesized for objects of different materials and masses without the detailed physical specification of an object that other synthesis methods require. This model enabled the generation of realistic experimental stimuli while also enabling us to manipulate the physical and statistical regularities present in impact sounds, which we leveraged to investigate their effect on perception.

To assess the basis of human perception we then asked humans to make physical inferences from the sound of a ball striking a surface. We found that listeners could accurately judge physical variables from sound. However, when both objects varied, as when comparing the mass of balls dropped on two different surfaces, inferences were impaired if the statistics of object resonances were manipulated to deviate from empirically measured physical regularities, with the extent of impairment depending on the extent of the deviation. The results indicate that humans have internalized knowledge of the distribution of object resonances as well as an intuitive understanding of how two objects interact to produce sound, and use this knowledge to perform physical perceptual inference. This internalized knowledge enables the perceptual separation of the different physical causes that combine to produce a single sound.

## Results

To study the perception of sound produced by physical interactions, we first sought to characterize how physical variables (mass, material etc.) affect sound. This characterization had two purposes: first, to aid in understanding the physical regularities that might be leveraged by our perceptual systems, in order to better enable us to formulate hypotheses, and second, to aid in synthesizing and manipulating sounds for experiments probing human perception.

Our general goal was to account for sound in terms of large-scale object properties that are relevant to everyday human experience, as these seemed likely to be targets for perceptual inference, and to be the basis of any implicit physical knowledge underlying perceptual inferences. Our approach builds on a large body of prior acoustics theory, but differs substantially from contemporary techniques which synthesize the sound made by individual objects using fine-grained physical descriptors of the object and computationally intensive simulations of acoustics^8–11^, and which are less well suited to our scientific purposes (in which it is useful to be able to generate large numbers of realistic object sounds without precisely specifying the object). Our approach also differs from classical techniques for deriving acoustic properties of objects, in that we augment physical theory with a statistical characterization of empirical measurements from actual objects. We emphasize that the purpose of our generative model was to enable physics-related perceptual science, not to produce maximally realistic sounds (for which a more complicated approach might have worked better).

### Physics-based synthesis model for impact sounds

The basis of our approach is the fact that in many situations, objects are well approximated as linear systems^6^ (Fig. 1). The vibrational response of an object can thus be modeled by the convolution of its impulse response and a time-varying force, as is created when two objects collide and then bounce apart. An impulse response is defined by the point of contact (where the force is applied) and the point of sound measurement. This point of measurement can be on the object surface, with each point on the surface having a distinct response. The point of measurement could also be some distance away from the object, in which case the response is a linear combination of the responses of each point on the object surface, with sound radiating into the surrounding medium from all points on the surface. In any real-world setting this externally defined impulse response would also contain the effects of sound reflections off of environmental surfaces. The impulse response is a useful characterization because it can be used to predict the sound at the point of measurement for any force at the point of contact.

At least two objects are involved in an impact event, each of which has a vibrational impulse response, and which together determine the force exerted at the point of contact. Consider the situation depicted in Fig. 1a, in which a ball is dropped on a surface, and for now, consider the impulse responses to be defined with respect to a point of measurement somewhere in the surrounding medium (as if a microphone was positioned near the objects). In the simplest case with a single impact (no subsequent bounces), the sound can be modelled as:

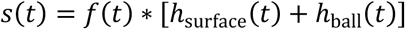

where *s* is the impact sound, *f* is the contact force, ℎ_surface_ is the impulse response of the impacted surface, and ℎ_ball_ is the impulse response of the ball. However, if the ball is small and dense, it will radiate much less sound than the surface^23,24,35^, and the contribution of its impulse response can be neglected (Fig. 1b). This simplification yields the following generative formulation for an impact sound:

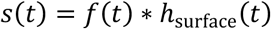

This formulation factorizes the acoustic waveform into a single impulse response and a time-varying contact force *f*(*t*) that depends on both objects and their momentum at impact (Fig. 1b). We adopt this problem setting to study the perceptual inference of physical variables from sound, as it retains the ill-posedness of the general problem, while reducing the number of factors to consider and manipulate. Fig 1c depicts the families of hypotheses we wish to distinguish in this problem setting. One class of hypothesis holds that humans derive acoustic cues from an impact sound and use those to judge physical variables. The other class of hypothesis holds that humans jointly infer the constituent causes of a sound, using knowledge of the distribution of impulse responses and contact forces along with their dependence on physical variables. The second class of hypothesis does not preclude the use of acoustic cues, but posits that any such cues are interpreted in light of the distribution of impulse responses and contact forces.

In this paper we sought to distinguish these two classes of hypotheses by asking whether the ability to infer one physical variable (e.g. ball mass) is influenced by the ability to infer another concurrent variable (e.g. surface material). We manipulated the ability to infer the concurrent variable by altering the extent to which one object’s impulse response conformed to the distribution of real-world impulse responses. In the sections that follow we characterize the distribution of everyday object impulse responses using large-scale acoustic measurements, use the measurements to build a generative model of object impulse responses, derive and validate a formulation of impact force, and assemble a generative model of impact sounds. We then use the generative model to study the human perception of impact sounds.

### Object impulse response measurement

Object impulse responses can be derived for simple geometric shapes^6^ or can be simulated with high-resolution physical models^9^. However, these simulations are computationally expensive and depend sensitively on precise knowledge of an object’s shape and internal geometry^9,36^. To avoid reliance on such techniques, we instead measured object impulse responses empirically from a wide range of everyday objects. Measurements were made by broadcasting pseudorandom noise into an object and re-recording the noise from the object’s surface, analogous to methods often used to measure the impulse responses of reverberant spaces^33^. We chose to measure impulse responses from the object surface in order to be able to make measurements in everyday environments (for which recordings with an external microphone would be contaminated by contributions from environmental reverberation). Contact speakers and microphones attached to the object’s surface were used to transmit vibrations into the object surface and record the result, from which the impulse response could be estimated (Fig. 2a; see Methods).

**Figure 2.**
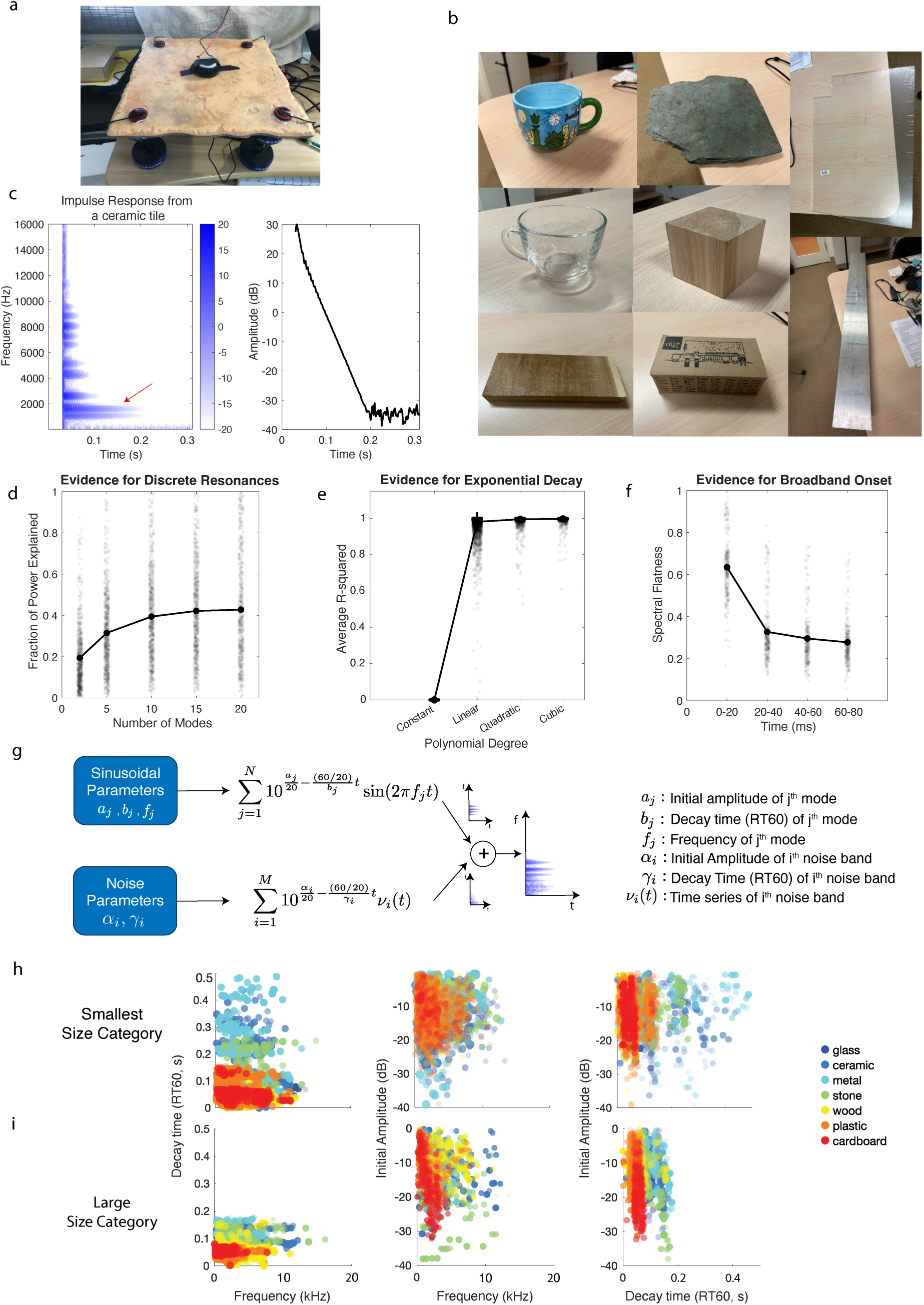
Object impulse responses. **a**. Apparatus for measuring object impulse responses. A noise signal is broadcast into an object with a contact speaker (in center of tile in photo), and re-recorded from the object’s surface via contact microphones (in corners of tile in photo). The object’s impulse response is estimated from the cross-correlation of the original noise signal and its re-recording. **b**. Example objects from impulse response survey. **c**. Spectrogram of example object impulse response (left; color scale denotes amplitude in dB) and the energy decay of one of its modes (right). **d**. Evidence for discrete resonant frequencies. Power accounted for by strongest modes of each impulse response. The contribution of sinusoidal modes to the impulse response can be accounted for by a fairly modest number of modes. Dots plot results for individual object impulse responses (horizontal position of each dot is randomly jittered for ease of visibility). **e**. Evidence of exponential decay of impulse response modes. Graph plots variance in the log-energy vs. time of each mode of each impulse response explained by polynomials of different orders. Analysis was restricted to the energy above the estimated noise floor. Nearly all the variance is accounted for by a linear function, indicative of exponential decay. Dots plot results for individual impulse responses. **f**. Evidence for broadband onset of impulse response. Graph plots spectral flatness of the power spectrum computed from successive 20-ms excerpts of impulse responses. High values of spectral flatness indicate a broadband spectrum. Dots plot results for individual impulse responses. **g**. Parametric model of object impulse response. **h**. Mode parameters for all objects in the Small size category (10-30 cm). Each dot plots a single mode from a single impulse response, color-coded by the object’s material. **i**. Mode parameters for all objects in the fourth size category (Large; 1-3 m). Same conventions as h.

To select objects that span the range of acoustic variability that humans encounter in daily life, we used an online survey. Participants reported the objects for which they had recently heard collision sounds, with separate prompts for each of five size categories ranging from less than 10 cm to greater than 3m in linear dimension. We combined their responses into a target list of objects (Table 1), and recorded as many of these commonly identified objects as was practical (see Methods and Table 2). The resulting set of objects spanned a wide range of material properties, sizes, and shapes (Fig. 2b). To our knowledge these measurements constitute the first large-scale characterization of everyday object acoustics.

**Table 1.** Material and size categories identified by the impact sound survey. The numbers listed indicate: No. of mentions (No. of unique objects; No. of objects that account for 50% of mentions). A small fraction of unique objects accounted for more than half of all responses. We only measured impulse responses from the four smallest size categories as our measurement method could not supply enough power to make useful measurements from the largest size category.

|  | <10cm | 10-30cm | 30-100cm | 1-3m | >3m |
| --- | --- | --- | --- | --- | --- |
| Metal<br>655 (125; 19) | 143 (20; 2) | 128 (22; 3) | 115 (35; 8) | 146 (25; 4) | 123 (23; 2) |
| Wood<br>609 (80; 11) | 35 (5; 1) | 54 (12; 2) | 113 (22; 3) | 245 (21; 2) | 162 (20; 3) |
| Composite<br>574 (177; 21) | 57 (24; 5) | 190 (42; 4) | 193 (42; 3) | 73 (39; 5) | 61 (20; 4) |
| Plastic<br>444 (89; 14) | 140 (22; 3) | 137 (21; 4) | 107 (24; 3) | 42 (11; 2) | 18 (11; 2) |
| Glass<br>142 (34; 11) | 13 (7; 3) | 28 (4; 1) | 17 (8; 3) | 61 (8; 2) | 23 (7; 2) |
| Ceramic<br>74 (14; 7) | 12 (2; 1) | 38 (4; 2) | 12 (4; 2) | 12 (4; 2) |  |
| Stone<br>54 (8; 4) | 6 (2; 1) | 4 (3; 1) |  | 11 (3; 1) | 33 (3; 1) |
| Rubber<br>51 (12; 3) | 24 (6; 1) | 16 (4; 1) | 11 (2; 1) |  |  |
| Paper<br>47 (12; 5) | 13 (5; 2) | 27 (5; 2) | 17 (2; 1) |  |  |
| Concrete<br>41 (3; 1) |  |  |  |  | 41 (3; 1) |
| Cardboard<br>24 (5; 3) |  | 7 (2; 1) | 14 (2; 1) | 3 (1; 1) |  |
| Plaster (wall)<br>24 (2; 1) |  |  |  |  | 24 (2; 1) |
| Drywall (wall)<br>19 (2; 1) |  |  |  |  | 19 (2; 1) |
| Brick (wall/road)<br>15 (2; 1) |  |  |  |  | 15 (2; 1) |
| Foam<br>10 (4; 2) | 3 (2; 1) | 7 (2; 1) |  |  |  |
| Asphalt (road) 8 (1;1) |  |  |  |  | 8 (1; 1) |
| Linoleum/Vinyl (floor)<br>8 (1; 1) |  |  |  |  | 8 (1; 1) |
| Laminate<br>(floor/counter)<br>3 (2; 1) |  |  |  |  | 3 (2; 1) |
| **Total**<br>**2802**<br>**(475; 107)** | **436**<br>**(95; 20)** | **636**<br>**(121; 20)** | **571 (141; 25)** | **593 (108; 19)** | **538 (97; 21)** |

**Table 2.** Material and sizes of all objects from which we measured and analyzed impulse responses.

|  | <10cm | 10-30cm | 30-100cm | 1-3m | Total (410) |
| --- | --- | --- | --- | --- | --- |
| Glass | 10 | 8 | 5 | 3 | 26 |
| Ceramic | 4 | 23 | 6 | 1 | 34 |
| Metal | 8 | 36 | 3 | 5 | 52 |
| Stone | 1 | 4 | 2 | 1 | 8 |
| Wood | 18 | 30 | 33 | 28 | 109 |
| Plastic | 69 | 70 | 18 | 5 | 162 |
| Cardboard | 1 | 12 | 2 | 4 | 19 |
| **Total (410)** | **111** | **183** | **69** | **47** |  |

### Real-world object measurements reveal stereotyped form of object impulse responses

Despite the wide variety of measured objects, there was considerable consistency in the form of the impulse responses. The example shown in Fig. 2c (for the object shown in Fig. 2a) illustrates the key properties when plotted as a spectrogram. First, consistent with classical theory, nearly all impulse responses exhibited a set of relatively discrete frequencies – the resonant modes of the object – evident as horizontal stripes in the spectrogram. The mode frequencies are inharmonic, unlike some other natural sounds (e.g. a plucked string, or many vocal sounds, in which the constituent frequencies are harmonically related, being integer multiples of a common fundamental frequency)^37^. Second, the power in these modes decays over time with a form that is closely approximated by exponential decay (Fig. 2c, right panel). Third, the impulse response typically begins with a broadband onset (i.e. a ‘click’) which decays rapidly, evident as the vertical stripe at the beginning of the spectrogram. This confirms the theoretical expectation that an impulse initially excites all frequencies, most of which decay very rapidly, leaving energy only in the resonant modes^7,38^.

We substantiated these observations quantitatively in the full set of measured impulse responses. First, we measured the fraction of the impulse response power that was accounted for by the N strongest spectral components (Fig. 2d). This analysis revealed that most of the power contributed by sinusoidal modes is due to the 10 strongest modes, illustrating that most impulse responses are dominated by a small set of relatively discrete modes. This analysis also revealed that the strength of the resonant modes (relative to the rest of the energy in the impulse response, which we will model with noise in the next section) varied considerably across objects. Second, we fit polynomials to the log-energy decay in these 10 modes, and found that most of the variance in the decay (excluding the noise floor) was accounted for by a linear function (i.e. an exponential function on a log scale), demonstrating that the modes nearly always decay exponentially (Fig. 2e). This result conveniently allows each mode to be summarized with three parameters: the frequency, the decay time, and the starting amplitude. Third, we measured the spectral flatness of the impulse responses in short time windows (Fig. 2f). The initial time window consistently had a more broadband spectrum than later windows, indicating the consistent presence of a broadband onset to the impulse response. These findings are consistent with classical theory^3–5^ and smaller scale measurements of object sounds^7^, but show that the theory holds broadly for everyday objects, and that the acoustic properties of everyday object have a fairly stereotyped form as a result.

### Parametric model of object impulse responses

The stereotyped form of object impulse responses can be captured with a parametric model (Fig. 2g) that sums sinusoidal resonant modes and decaying noise bands (critical for correctly capturing the onset):

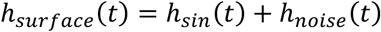

The sinusoidal component is given by:

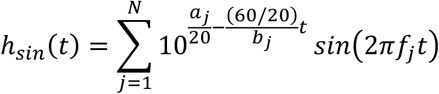

where a_j_ is the initial amplitude, b_j_ is the decay time (RT60; the time for the amplitude to decay by 60 dB) and f_j_ is the frequency of the j^th^ mode.

The noise component is given by:

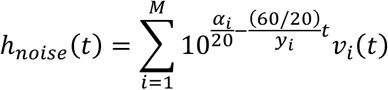

i.e., a sum of noise bands *v*_*i*_ that tile the frequency spectrum with fixed cutoffs, parameterized by a starting amplitude α_i_ and decay time (RT60) y_i_ for each noise band.

We fit the parameters of this model to each measured object impulse response, using gradient-based optimization (see Methods). This yielded a description of each impulse response in terms of its mode frequencies, decay rates, starting amplitudes, and noise parameters.

### Different impulse responses from an object are similar

For each object we measured multiple impulse responses with different surface microphone locations. Each of these impulse responses summarizes the sound that would radiate from a point on the object’s surface given a force applied at the speaker location. Visual inspection of impulse responses from the same object reveals a fair degree of consistency along with some variation (Supplementary Fig. 1a). To quantify this consistency, we first measured the difference in the average mode parameter values for all pairs of impulse responses within an object, and compared it to pairs from different objects (Supplementary Fig. 1b). This analysis revealed that mode parameters were substantially more similar for impulse responses from the same object than for impulse responses from different objects. We then assessed the extent to which the different impulse responses from the same object share the same mode frequencies. This analysis showed that pairs of impulse responses from the same object tend to share a considerable number of modes (6.3 on average, SD = 0.91; Supplementary Fig. 1c), and that there are very few modes that are only present in one of the impulse responses measured from the same object (Supplementary Fig. 1d). Given the reversibility of linear systems, the same conclusions hold for impulse responses from different points of contact (as if we had measured impulse responses from one point but with different speaker locations).

These results confirm that real-world objects possess a modest set of resonant modes, and that any single impulse response measured from the object surface tends to include a subset of them. This finding suggests that a single impulse response is likely to be representative of the modal properties of an object, and thus is also likely to be representative of the impulse response for an external point of measurement that would combine the sound from different points on an object’s surface (discounting effects of environmental reverberation that will also contribute to such an impulse response in natural conditions).

### Object impulse response statistics – effects of material and size

It was also apparent from examination of the impulse responses that there were pronounced effects of material, and to a lesser extent, object size. To illustrate these effects we plot the mode parameters for each resonant mode of each of the objects we measured in the Small (10-30 cm) and Large (1-3 m) size categories (Fig. 2g&h), color-coded by the object material (see Supplementary Figs. 2-4 for plots for all materials and size categories). These plots show consistent variation in mode properties with material (harder materials like metal and ceramic show longer decay rates and higher frequency modes, consistent with prior theory^14–16,18,39^). There is also some variation with size (larger objects tend to have lower frequency modes, consistent with prior theory, and shorter decay times, plausibly due to a tendency for larger objects to be externally damped to a greater extent than smaller objects; Supplementary Fig. 5). The effects of material were fairly consistent across object sizes (Fig. 2g & h; Supplementary Fig. 2), and the effects of size were fairly consistent across material (Supplementary Figs. 3-5), although there is variation that may reflect particular shapes being most common in certain size/material combinations (e.g. mugs being the most common small ceramic object, and tiles being the most common large ceramic object; see Discussion). It is also apparent that although there is variation in average mode properties as a function of material and size, the distributions for different materials and sizes overlap substantially.

### Generative model of impulse responses

We used these measurements to build a generative model of impulse responses (Fig. 3a). We fit multivariate Gaussian distributions to the three parameters of a mode (frequency, starting amplitude, and decay time) and the two parameters of the noise bands (starting amplitude, decay time) conditioned on material and size. To generate an impulse response, we sampled each of 10 sinusoidal modes independently from the same 3D Gaussian distribution. Because the noise bands had fixed frequencies, we used a different 2D Gaussian for each band, from which we sampled the amplitude and decay time.

**Figure 3.**
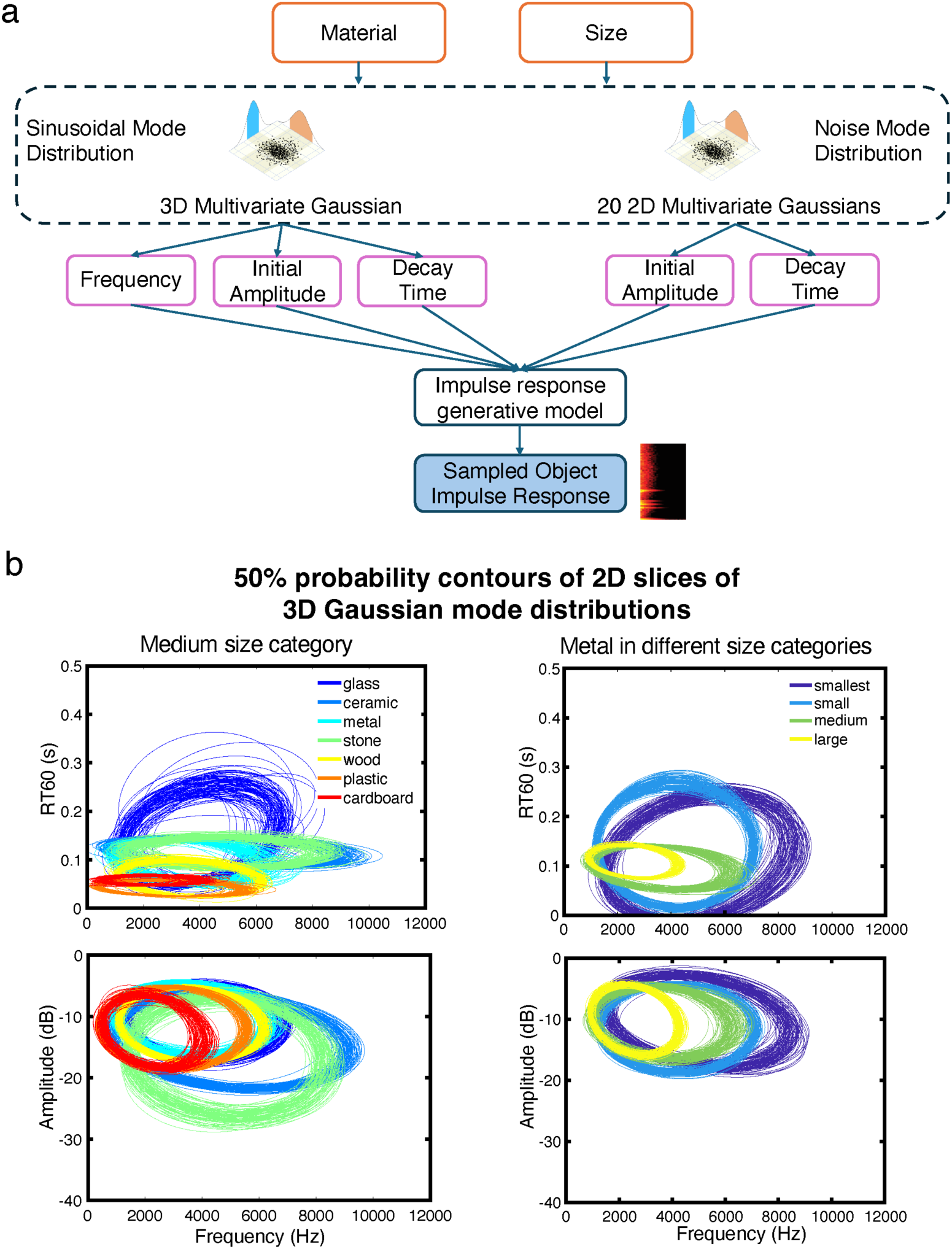
Generative model of impulse responses. **a**. Mode parameter dependencies were modeled with multivariate Gaussian distributions conditioned on the object material and size. **b**. Constant probability ellipsoids (half-maximum of probability density) from fitted Gaussian distributions. To provide a sense of the stability of the covariance between parameters, we have plotted ellipsoids from Gaussians fit to different bootstrap samples of the impulse responses for each material and size category. Note that Gaussian distributions were truncated at zero for all dimensions. Harder materials (glass, ceramic, metal and stone) tend to have modes with higher frequencies and longer decay times. Larger objects tend to have lower frequency modes. Both results are consistent with classical theory but show that it holds for real-world objects. Object size ranges were <10cm (Smallest), 10-30cm (Small), 30-100cm (Medium), and 1-3m (Large).

The fitted distributions reflect the effects of material and size evident in the scatter plots of Fig. 2g&h, but also reveal covariance between the three mode parameters, evident in constant-probability ellipsoids that are oriented in some cases (Fig. 3b). Supplementary Fig. 6 shows the distribution parameters for the noise bands (the amplitude of the noise bands is higher in softer materials, as expected given that these materials have less pronounced resonances).

The generative model can be used to sample impulse responses as part of the larger model of impact sounds described next. It can also be used to measure the likelihood of an impulse response under the real-world distribution, which we will use later in the paper to make predictions about the effect of alternative generative models on human perception.

### Physics and acoustics of impact force

The other essential ingredient in impact sound generation is the force created as objects come into contact, compress against each other, and then push each other apart. We compute contact force from an extension of a classical spring model^40^. The spring model assumes that the force between a ball and the surface it impacts depends linearly upon the distance by which they are compressed when in contact. This formulation yields a contact force as a half-cycle of a sinusoid, with the spring constant determined by the mass of the impacting object and the stiffnesses of both objects. The typical duration of such a force is less than a millisecond. As such, the primary acoustic effect of the contact force is to filter the object impulse response, with the following transfer function:

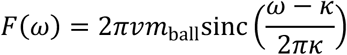

where *ω* is frequency, 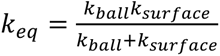 is the equivalent spring constant (i.e. resistance to compression) of the ball and surface, *v* is the velocity of the ball at impact, *m*_ball_ is the mass of the ball, and 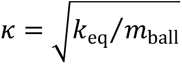. The overall amplitude of this transfer function (and thus the resulting sound) increases with the ball mass, as expected. Moreover, the function peaks at a frequency of *κ*⁄(2*π*). The peak frequency thus decreases for heavy balls and/or softer materials, as both of these increase the time the surface and ball are in contact, i.e., the width of the half-cycle sinusoid, resulting in a more low-pass filter (Fig. 4a&b). The formulation thus encompasses the two main acoustic cues previously associated with mass (amplitude and spectral centroid)^23,24^.

**Figure 4.**
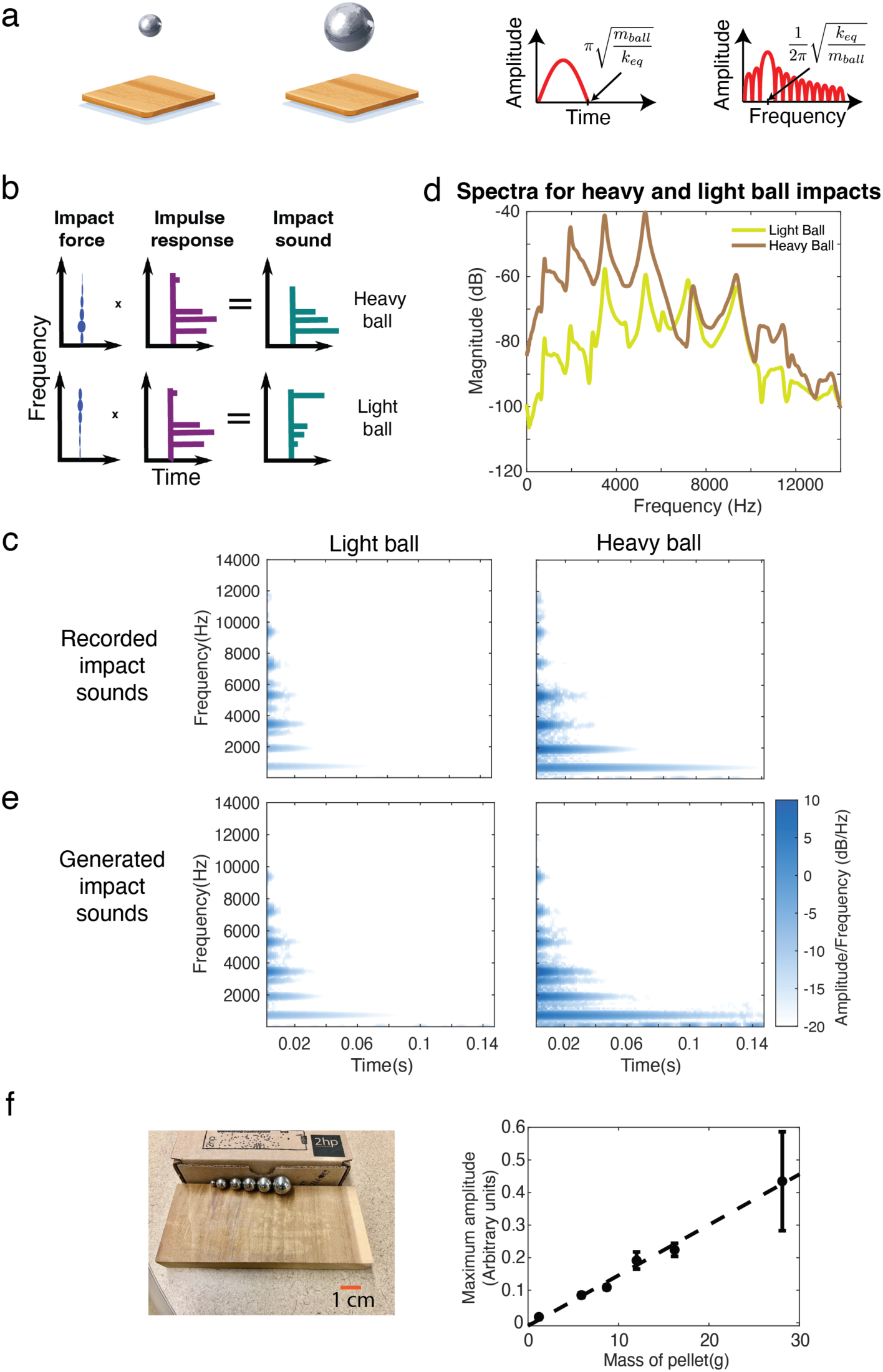
Effects of impact force. **a**. Acoustic effects of impact force. The force produced by an impact increases and then decreases over time as the objects come into contact, compress, and then move apart. The time-varying force predicted by our model is shown for two different masses. In both cases the forcing function is approximately a half-cycle of a sinusoid (here the masses are small enough that the force stays in the linear regime). **b**. Schematic depiction of effect of impacting object mass on the resulting impact sound. The peak of the transfer function varies with mass, filtering the impacted object impulse response to produce an impact sound. **c**. Spectrograms of recorded impact sounds from different impacting object masses. The heavier mass causes the lower frequency modes of the surface to be accentuated. **d**. Power spectra of the recorded impact sounds shown in d (shown to make the mode amplitudes more easily visible). **e**. Spectrograms of synthesized impact sounds, generated from a simulated contact force for two different masses, using the impulse response measured from the object shown in b. The generative model successfully captures the effect of mass on the impact sound. **f**. The maximum amplitude of recorded impact sounds produced by metal pellets of different masses. Small masses dropped from modest heights produce forces that remain in the linear regime, and that increase approximately linearly with mass.

We modified the spring model to account for limits on linearity that occur in real-world objects, which have limits on the extent to which they can compress (see Methods), but the qualitative effect of the contact force remains, with heavier masses and softer materials accentuating lower frequencies of the impulse response. To generate the force for an impact sound, we set the object stiffnesses to material- and size-specific values (see Methods; note also that for some experiments we held stiffness constant to isolate effects of the impulse response statistics on perception).

We validated the theoretical predictions for the effects of object mass using recorded impact sounds generated by dropping pellets of different masses onto object surfaces. This comparison showed the expected effects of object mass, with heavier objects preferentially exciting lower frequency modes (Fig. 4c&d). If our simulated contact force is convolved with the measured impulse response for an object, the resulting sound is similar to the recorded audio from an actual impact in the world (Fig. 4e), helping to validate the model. The recordings also help validate the effect of mass on amplitude: sound amplitude increased roughly linearly with mass for small pellets that stay within the linear regime of the forcing function (Fig. 4f), consistent with previous such measurements^23^.

### Generative model of impact sounds

By combining the models of impulse responses and contact forces, we obtain a generative model of impact sounds (Fig. 5a). The model generates a sound from a small set of physical variables: the object mass, material, size, and speed of impact, providing a tool we can use to synthesize stimuli and study perception.

**Figure 5.**
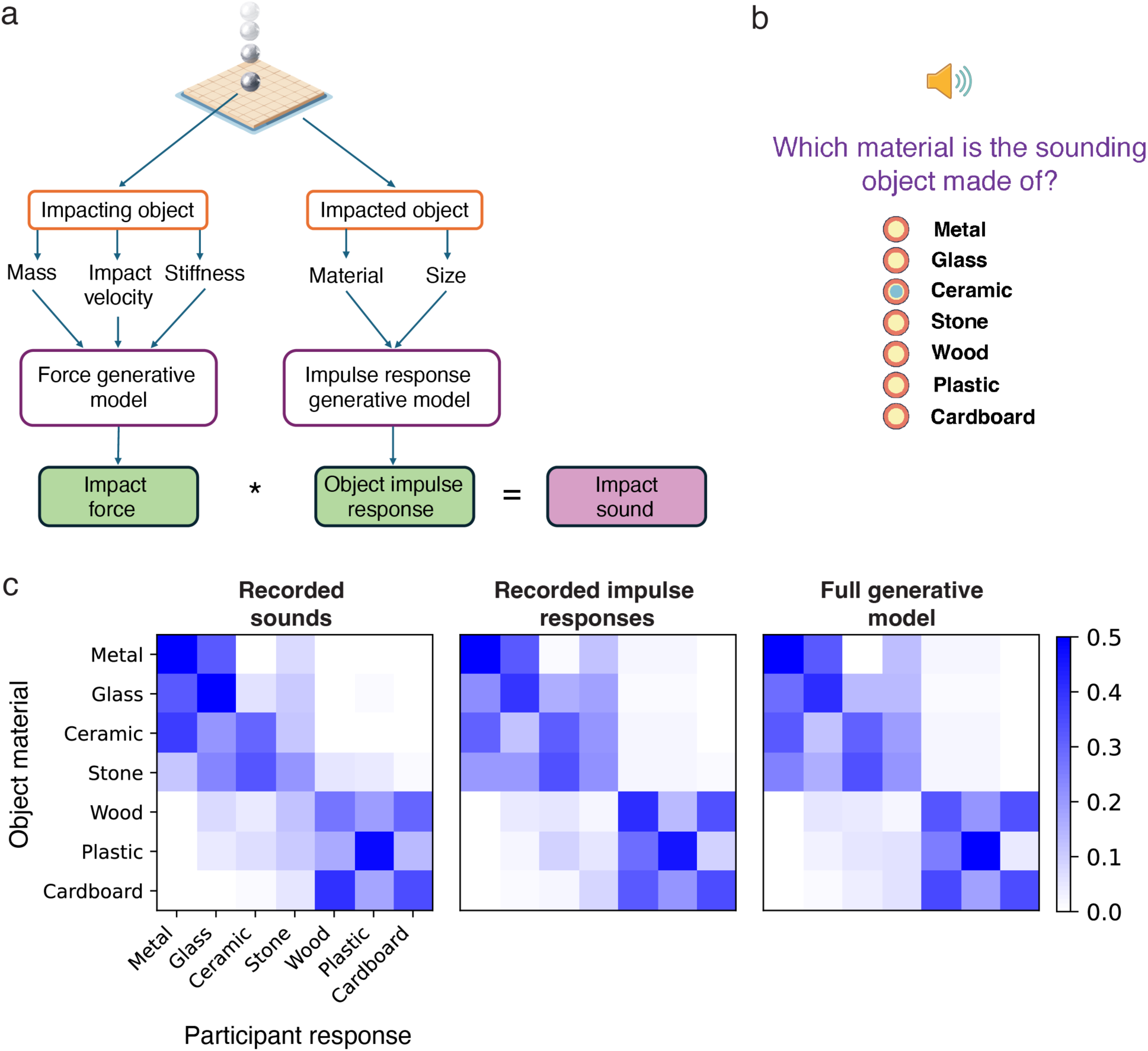
Generative model of impact sounds and its validation. **a**. Generative model of impact sounds. The full generative model combined the generative models for impact sounds and contact force shown in Figs. 3 and 4. **b**. Experimental task for Experiment 0. Listeners heard a sound and selected the object material from a list of seven choices. The sound was either an audio recording of an impact sound produced by a small pellet dropped on an object, a synthetic sound generated from a recorded impulse response, or a synthetic sound generated from a synthetic impulse response sampled from the full generative model. **c**. Confusion matrices for sounds from each experimental condition. We note that this experiment included only two of the four size categories, and used material-specific stiffness values in the synthetic sounds, and it is thus unsurprising that the pattern of confusions is somewhat different from that measured in Experiment 2.

### Experiment 0: Validation of impulse response recordings and impact sound synthesis

Because our approach relied on recorded impulse responses, and on impact sounds synthesized from a generative model, it was important to confirm that these were sufficient to render relatively realistic sounds. We validated the resulting synthetic sounds with an experiment measuring material identification using recorded and synthetic sounds. There were two reasons for using a material recognition task. First, material recognition yields a relatively high-dimensional result (a confusion matrix), providing a strong test of whether synthetic sounds capture the perceptual effects associated with real-world sounds. Second, material recognition seemed more likely to be robust to uninteresting differences between recorded and synthetic sounds, such as background noise or reverberation that would inevitably be present to some extent in recorded but not synthetic sounds.

Participants heard a single impact sound and categorized the material into one of seven categories (Fig. 5b). We compared material recognition for recorded impact sounds, sounds rendered from the generative model, and sounds rendered from the generative model using recorded impulse responses instead of sampled impulse responses. The latter two conditions used material- and size-specific values of stiffness, such that both the impulse response and forcing function contained information about material (as in real-world sounds). We used objects from the middle two size categories (10-30 cm and 30-100 cm), as these were most amenable to generating clean impacts (see Methods).

As shown in Fig. 5c, humans were well above chance at identifying an object’s material from sound, and performed similarly across the three conditions (recorded sounds: 38.5% correct; sound synthesized from recorded impulse responses: 37.7% correct; sounds synthesized from the full generative model: 38.2% correct). Humans nonetheless made many errors, consistent with prior studies^41^. These errors plausibly reflect the overlap between the distributions of impulse response properties of different materials, evident in Figs. 2, 3, and Supplementary Figs. 2-4. The confusion matrices were strongly correlated (recorded sounds vs. sounds synthesized from recorded impulse responses: r=0.94, p<.001; recorded sounds vs. sounds synthesized from the full generative model: r=0.95, p<.001). These results provide evidence that the assumptions underlying the generative model and impulse response recordings are sufficiently accurate as to produce synthetic sounds that are perceived in much the same way as real-world sounds.

### Lesioned variants of generative model

We investigated the perceptual importance of the various physical regularities built into the model using “lesioned” model variants that each violated some aspects of the full generative model: the covariance between parameters, the variance over individual parameters, the number of modes, the shape of the distributions, or the type of decay (Fig. 6a). These variants were chosen to produce impulse responses that deviated from the real-world impulse response distribution to varying extents, but were not otherwise particularly principled. We quantified this deviation by measuring the log-likelihood under the real-world distribution of impulse responses sampled from each condition (Fig. 6b). All of the model variants produced impulse responses that were less likely under the real-world distribution (compared to Recorded and Full model impulse responses), but the magnitude of this effect varied across conditions, as intended.

**Figure 6.**
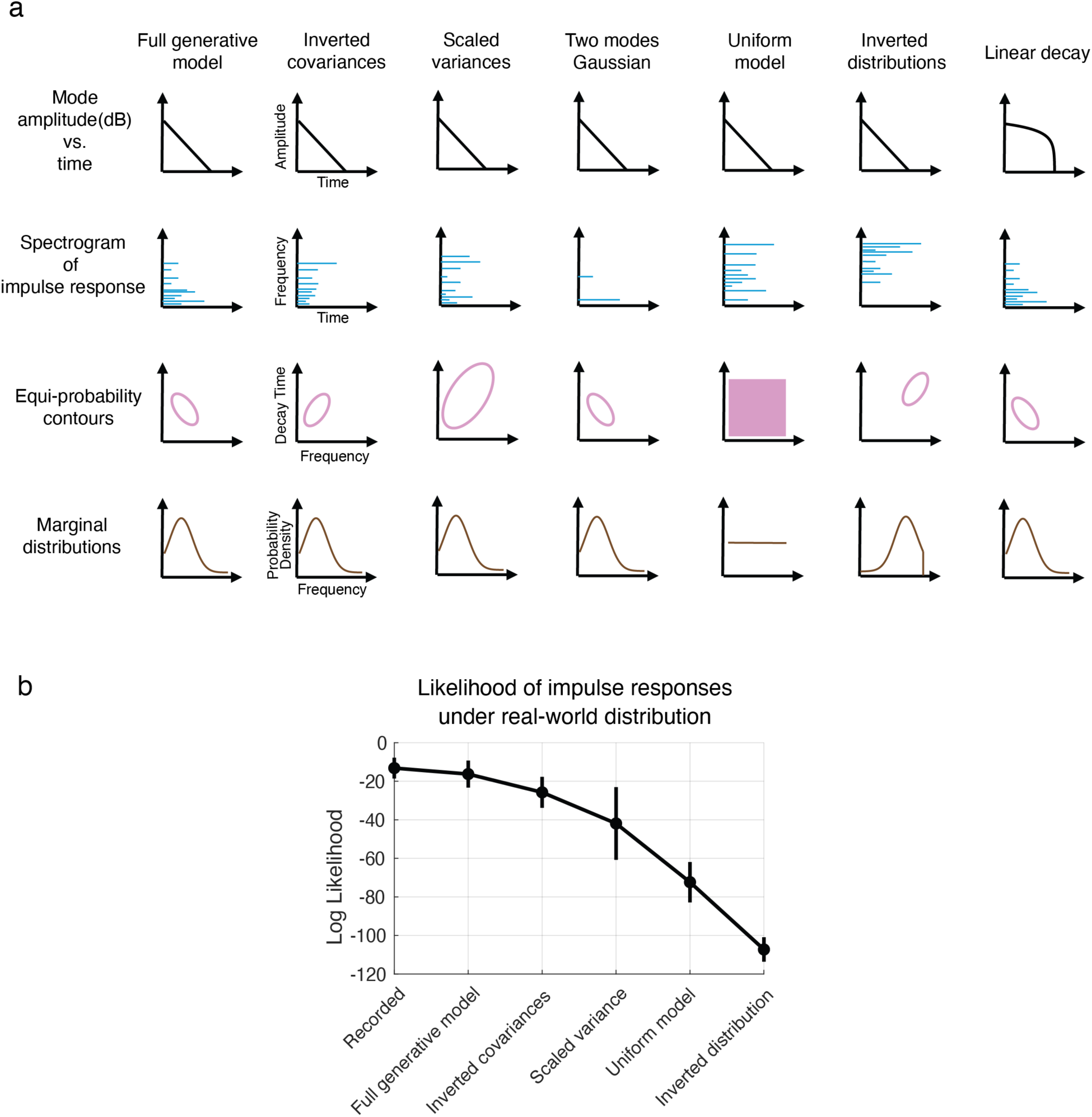
Lesioned variants of generative model. **a**. Schematic depiction of variants of generative model used in perceptual experiments. Top row shows the energy decay of an example sinusoidal mode, which was exponential in the full generative model (appearing linear on a dB scale) but that was altered to be linear in one of the model variants (appearing concave down on a dB scale). Second row shows schematic spectrograms of impulse responses. The full generative model produces impulse responses that have 10 modes, but this was reduced to 2 modes in one of the model variants. Third row shows equiprobability contours for the distributions from which mode parameters are sampled. In the full generative model these distributions are Gaussian, with covariances measured from real-world impulse responses. In two of the model variants the covariance is inverted, producing the opposite relationship between mode parameters (shown here for frequency and decay time, also evident in the schematic spectrogram); in one of these variants the variance is also scaled up. In two other model variants the distributions take a different form, with one being uniform and another being “inverted”, having high probability where the full model has low probability, and vice versa. Fourth row shows distributions marginalized over decay time and amplitude for ease of visualization. Note that Gaussian distributions were truncated at zero on the frequency axis. **b**. Log-likelihood of impulse responses from model variants, evaluated under the full generative model. We omitted the Two modes and Linear decay conditions as their mode parameters were taken from the full model.

### Experiment 1: Realism of synthesized impact sounds

We first tested whether the realism of sounds produced by our generative model depended on the physical and statistical regularities captured by the model. Participants heard two sounds and judged which was from a real object (Fig. 7a). One of the sounds was generated from a recorded impulse response, and the other was generated using the generative model or one of the model variants shown in Fig. 6a.

**Figure 7.**
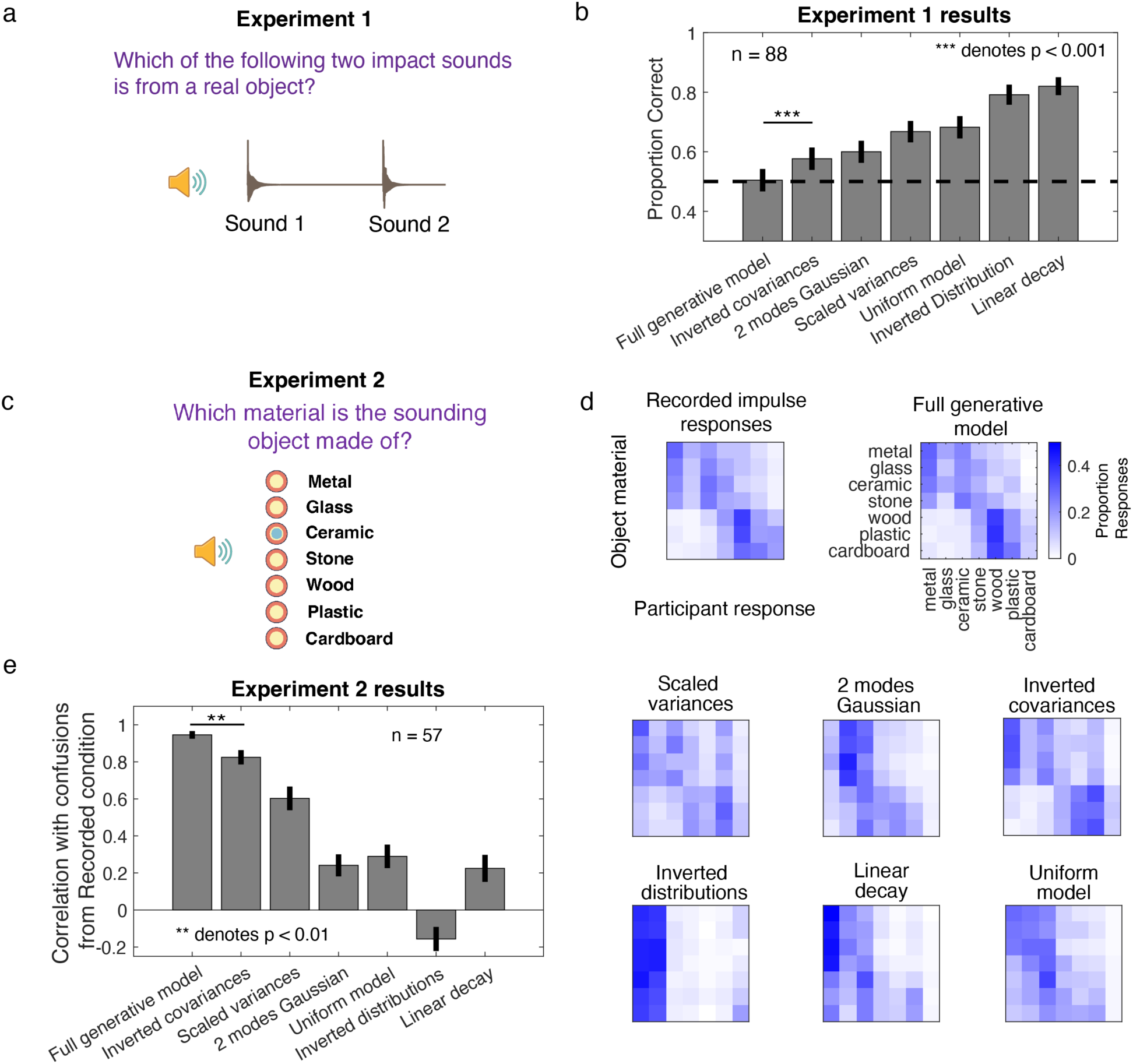
Realism of synthetic impact sounds depends on physical and statistical regularities. **a**. Experimental task for Experiment 1. Listeners heard two sounds and judged which was from a real object. On each trial, one sound was generated from a recorded impulse response, and the other was generated from a synthetic impulse response. The synthetic impulse response was either sampled from the generative model or from one of several “lesioned” model variants. **b**. Results of Experiment 1. Participants were unable to distinguish sounds synthesized from the generative model from those measured from actual objects, but each of the lesioned models was distinguishable to some extent. Error bars plot SEM, obtained via bootstrap. Dashed line plots chance performance level. **c**. Experimental task for Experiment 2. Listeners heard a sound and selected the object material from a list of seven choices. The sound was either generated from a recorded impulse response or from a synthetic impulse response sampled from the generative model or a lesioned version of the model. **d**. Confusion matrices for sounds from each experimental condition. We note that this experiment included all four size categories, and used a single stiffness value for all materials (to isolate the contribution of the impulse response on material perception, for ease of comparison with Experiments 3 and 4). It is thus unsurprising that the pattern of confusions is somewhat different from that measured in Experiment 0. **e**. Correlation between confusion matrix for sounds from recorded impulse responses and the matrix for each model condition. Error bars plot SEM, obtained via bootstrap.

As shown in Fig. 7b, human listeners were unable to distinguish sounds from the generative model from those made from recorded impulse responses (t(86) = 0.420, p = 0.68; d =0.046). However, sounds from the model variants (each of which violated a physical or statistical regularity found in real-world objects) were distinguishable from those from the Recorded condition. The size of this effect depended on the variant, but even the most subtly different condition (Inverted covariance) was significantly more distinguishable than the Full model (t(86) = 4.38, p < 0.001, statistically significant after Bonferroni correction; d >= 0.50). This result suggests that our model is successful in capturing the perceptually important features of real-world impacts, and that this success depends on its physically inspired structure and empirically captured statistical regularities.

### Experiment 2: Material recognition of synthesized impact sounds

As a second test of the perceptual significance of the physical and statistical regularities of the generative model, we tested the effect of model lesions on material perception. Participants heard a single impact sound and categorized the material into one of seven categories (same task as Experiment 0; Fig. 7c). The sound was generated using a recorded impulse response, using the generative model, or using one of the model variants. Here and in subsequent experiments, we used a single value for stiffness, so as to isolate the effect of the object impulse response on perception.

Because the confusion matrix provides a rich perceptual signature with which to probe human sensitivity to the acoustic structure of the model, our main analysis compared the confusions resulting from sounds from each model variant to that for sounds generated from recorded impulse responses. As shown in Fig. 7d, the confusions produced by actual object impulse responses were closely mirrored by those for samples from the full generative model. We note that the confusions differed somewhat from those of Experiment 0 (Fig. 5c); these differences are unsurprising given that Experiment 2 eliminated material cues from the forcing function and also sampled from all four size categories. More notably, confusions for each of the model variants deviated from the pattern seen for the full generative model. We summarized the similarity between confusion matrices with a correlation coefficient between the matrix for actual impulse results and that for each of the model-generated sounds. This correlation was close to 1 for sounds from the full generative model but was lower for each of the model variants (p<.003 for each paired comparison, via permutation test, significant after Bonferroni correction; difference in correlation >= 0.12). This result indicates that the generative model captures the acoustic regularities underlying human material recognition from sound, and that the lesions interfere with the human perception of material.

Overall, Experiments 1 and 2 confirm that humans are sensitive to the acoustic structure of impact sounds and associate this structure with object properties. The experiments also establish that our generative model captures much of the relevant acoustic structure.

### Mass discrimination within and across impacted surfaces

One of the most interesting, and most neglected, aspects of the perception of physical interactions is the question of whether and how humans can solve the ill-posed problems created by the confluence of physical variables from multiple objects. We investigated this question in two scenarios (Experiments 3 and 4, respectively). The first scenario was that of estimating the mass of a dropped object. This problem is in general ill-posed because the mass of the dropped object and the impulse response of the impacted surface both influence the sound that is produced (Figs. 1 and 4). To study this problem, we designed the task shown in Fig. 8a-c. In this scenario, a heavy ball and a light ball are dropped onto surfaces. The task is to determine which ball is heavier. We chose a discrimination task rather than an estimation task with a single impact sound because it was not obvious that typical human participants would be able to express well-calibrated absolute mass judgments^23^. We asked whether the ability to infer the mass of the ball depended on whether the impulse response of the impacted surface was drawn from the natural distribution (which influences the extent to which the material of the impacted surface can be correctly inferred, as shown in Experiment 2).

**Figure 8.**
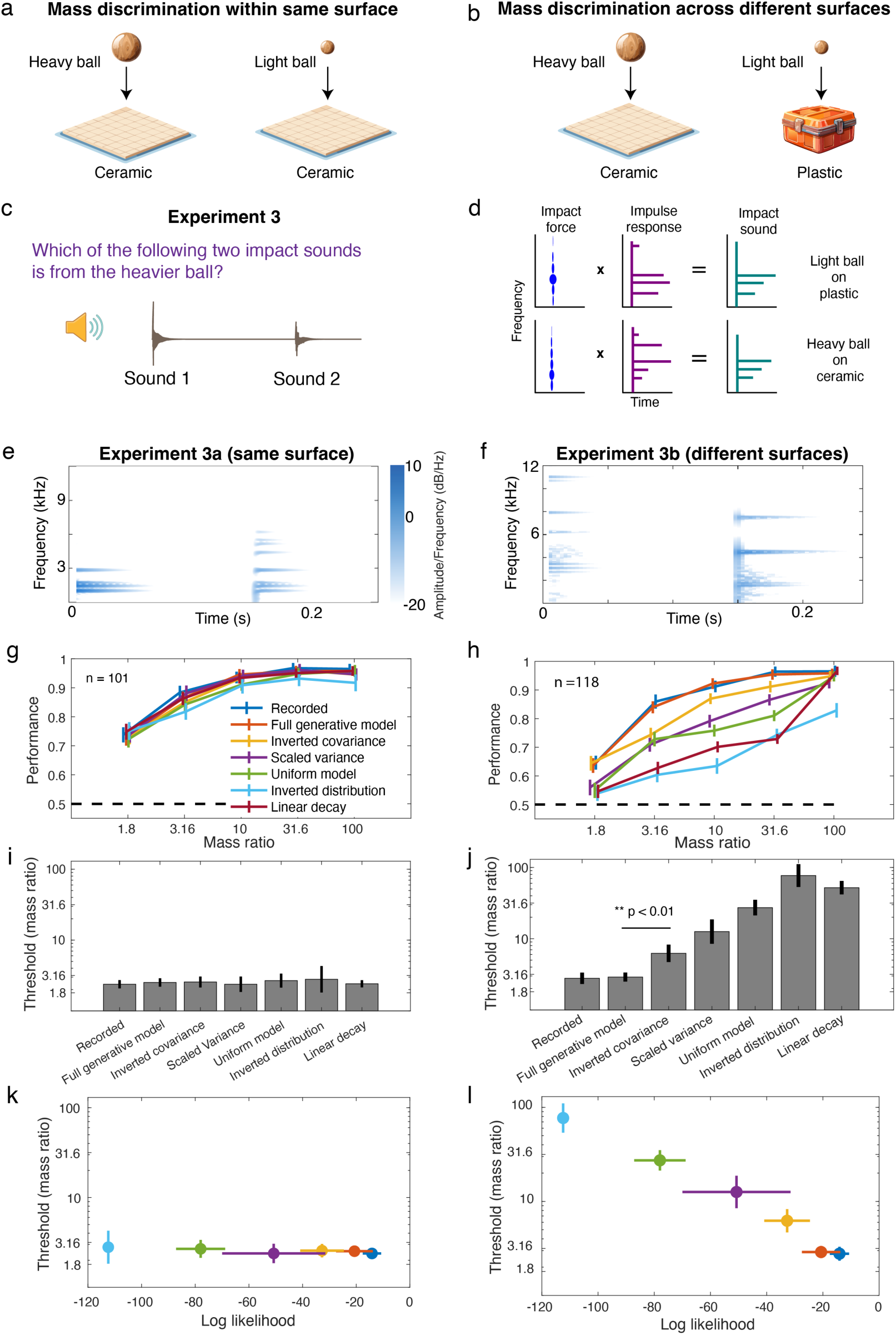
Mass discrimination within and across impacted surfaces. **a&b**. Scenarios explored in Experiment 3. Two balls of different masses are dropped on either the same object, or on two different objects. **c**. Task of Experiment 3. Listeners heard two impact sounds and judged which was produced by the heavier ball. The experiment manipulated the difference between the two masses and the extent to which the impact sounds were faithful to the physics-induced regularities of real-world impacts. **d**. The joint influence of ball mass and surface material on the spectrum of the impact sound renders the inference of mass ill-posed. **e**. Spectrogram of stimuli on example trial of Experiment 3a, in which both sounds are rendered from the same metal surface. The object resonances (modes) are the same in the two stimuli (note that the three prominent frequencies in the first stimulus are also present in the second). As a result, the discrimination can be performed using simple acoustic cues of level and centroid (note that the peak amplitude is higher in the first stimulus, and that the centroid of the spectrum is lower). **f**. Spectrogram of stimuli on example trial of Experiment 3b, in which one object is ceramic and the other is stone. Here the object resonances (modes) are different in the two stimuli (note that each stimulus contains a different set of frequencies). As a result, the simple acoustic cues that are diagnostic in Experiment 3a are not as diagnostic for solving the task. **g**. Proportion correct plotted as a function of the mass ratio between the two balls, for Experiment 3a. Error bars plot SEM, obtained via bootstrap. Dashed line plots chance performance level. **h**. Proportion correct plotted as a function of the mass ratio between the two balls, for Experiment 3b. **i&j**. Discrimination thresholds for Experiments 3a and 3b. Thresholds were defined as the mass ratio resulting in 80% correct, estimated by fitting splines to the data in d. Error bars plot SEM, obtained via bootstrap. **k&l**. Comparison of discrimination thresholds from Experiments 3a and 3b with log-likelihood of impulse responses. The linear decay condition is omitted as the impulse response parameters were taken from the Full model, and thus do not reflect the deviant features of that model variant. Error bars on thresholds plot SEM, derived via bootstrap. Error bars on log likelihood plot SD across impulse responses.

If the two balls are dropped on the same surface from the same height (Fig. 8a), mass discrimination can be straightforwardly performed by comparing the amplitude and frequency content of the resulting sound (because the heavier ball will produce a sound higher in amplitude and with more power at low frequencies; Fig. 4b-d). However, if the two surfaces are of different materials (Fig. 8b), the task becomes ill-posed. The generative components underlying two example impact sounds are shown in Fig. 8d. As discussed earlier, impact sound generation can be conceptualized as a reweighting of the object impulse response frequencies by the transfer function of the impact force. As a result, a heavier object dropped on a harder surface (e.g. ceramic) and a lighter object dropped on a softer surface (e.g. plastic) might be difficult to distinguish simply via the balance of high or low frequencies.

We hypothesized that listeners might overcome the ill-posed nature of this problem by leveraging implicit knowledge of impulse response regularities and the physics-induced structure of sound generation. For instance, knowing that harder objects tend to have modes that are both higher frequency and that have slower decay rates (Fig. 2h&i) might enable a listener to discount the contribution of the impulse response to the sound spectrum, enabling estimation of the mass. This hypothesis predicts that mass inference should be impaired if the impulse response of the struck surface deviates from the naturally occurring impulse response distribution, since in such a scenario a listener’s implicit knowledge of impulse responses would be less useful in factoring out the impulse response from the force. We tested this hypothesis by asking whether humans can perform this discrimination, and testing whether their performance was dependent on the stimuli obeying the regularities found in real-world impulse responses.

A priori it was not clear what to expect, in part because it was not clear that the impulse responses of real-world objects are sufficiently constrained that their distribution would be useful in inferring mass. Thus, it was not obvious that performance would depend on whether impulse responses were drawn from the Full generative model (approximating the real-world distribution) vs. one of the lesioned generative models.

### Experiment 3a: Mass discrimination within the same impacted surface

We first tested human listeners in an experiment in which the same impulse response was used for the two sounds, simulating the setting in which the two balls are dropped on the same surface (Experiment 3a; Fig. 8a). In this case the effects of the impulse response do not have to be separated from that of the impact force; mass can be discriminated simply by comparing the acoustic cues that are correlated with mass (e.g. amplitude and spectral centroid; Fig. 4). Moreover, the trial structure was intended to make these cues obvious to the listener (Fig. 8e): the two sounds on a trial contain the same frequencies (object resonances), such that there is a single factor of variation distinguishing the pairs of stimuli. If the different stimulus conditions (synthesized from different generative model variants) preserve these individual cues as intended, performance should be comparable for all conditions.

As shown in Fig. 8g, performance was indeed similar for all model variants. This result indicates that when the problem is not ill-posed, listeners can discriminate mass using cues that are preserved by all the model variants.

### Experiment 3b: Mass discrimination across different surfaces

We then tested human listeners in an experiment in which the two impact sounds were synthesized from different impulse responses, as if the balls were dropped on different surfaces, rendering the discrimination ill-posed (Experiment 3b; Fig. 8b&8d). Here the two stimuli on a trial have distinct object resonances (Fig. 8f). Unlike Experiment 3a, there is not a single acoustic dimension that clearly distinguishes the two stimuli on a trial. In this case we observed large differences between conditions (Fig. 8h). Performance improved as the simulated mass difference between the two balls increased, but was worse for the five conditions that violated real-world impulse response regularities.

We quantified performance as the threshold mass ratio required to yield 80% correct (Fig. 8i&j). These thresholds were similar across conditions for Experiment 3a (same object), but varied across conditions for Experiment 3b (different objects). In particular, the two conditions with realistic impulse responses yielded similar thresholds (p=0.77, via permutation test; ratio between thresholds = 1.04), but each of the five conditions from the alternative generative models yielded significantly worse performance (p<=.001 for each paired comparison with the Full model condition, statistically significant after Bonferroni correction; ratio between thresholds >= 2.15). Moreover, in the conditions in which the impulse response statistics were varied, the thresholds co-varied with the average log-likelihood of the impulse responses (Fig. 8k&l), consistent with the idea that humans make use of a prior over impulse responses to separate out the impulse response from the impact force (that contains information about mass). This result confirms the key prediction of the hypothesis that listeners overcome the ill-posed nature of the problem by leveraging implicit knowledge of the statistical and physical regularities of sound generation.

### Results not explained by individual acoustic cues

To further confirm that the differences in human performance across conditions could not be explained by differences in individual acoustic cues, we implemented two “cue discriminator” models that performed the task by comparing either the spectral centroid or amplitude of the two sounds, and picking the lower or higher amplitude sound, respectively. These two cue discriminator models performed similarly across conditions (Fig. 9a-c), as expected. Moreover, the centroid discriminator underperformed humans for the two naturalistic conditions (Recorded and Full model; Fig. 9c). These results indicate that all stimulus conditions contain similar low-level acoustic cues, as intended, and underscore that humans are not performing the task with a simple comparison of these cues.

**Figure 9.**
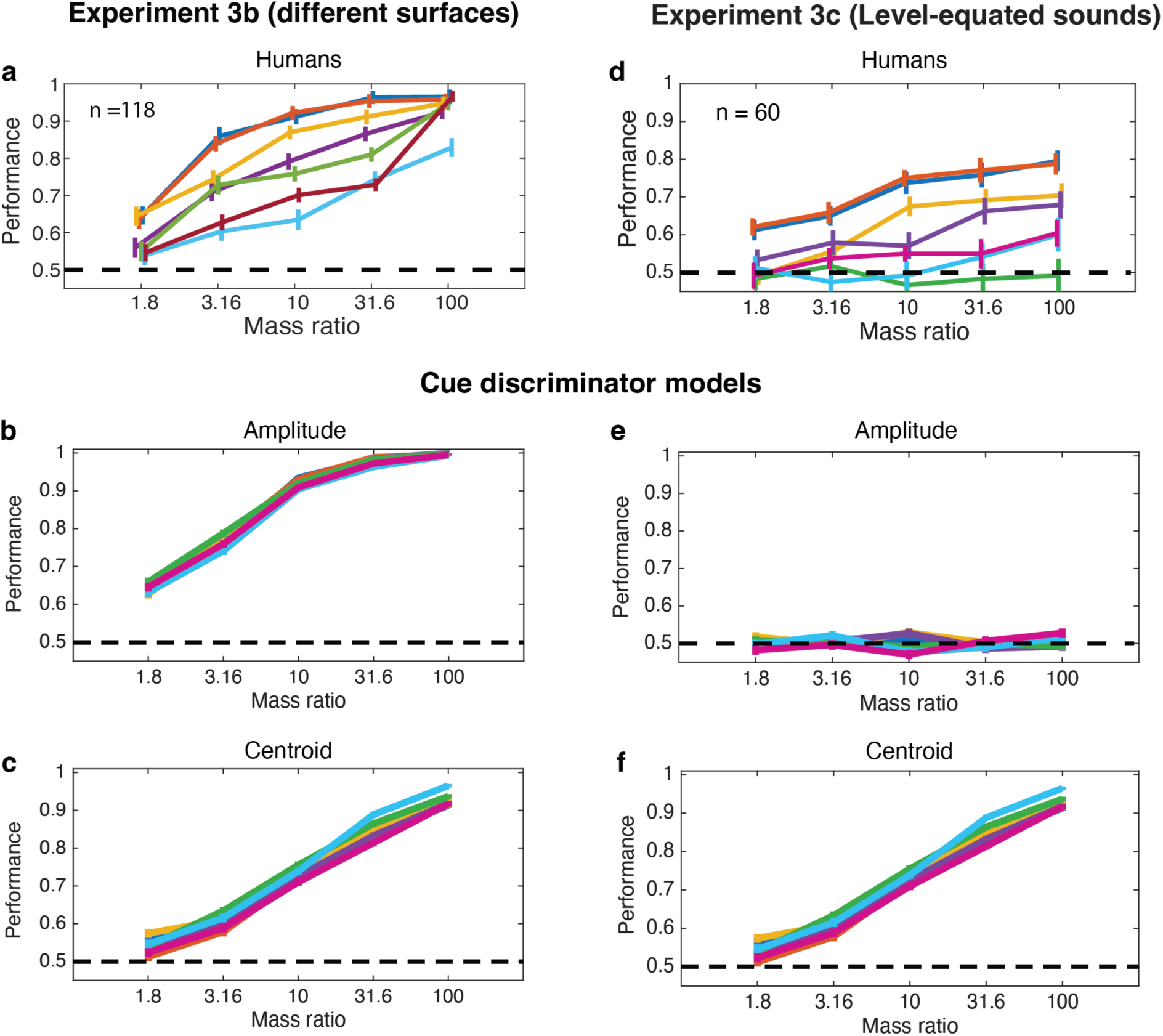
Human mass discrimination is not explained by single acoustic cues. **a**. Results of Experiment 3b, replotted for comparison with other panels. Here and elsewhere, error bars plot SEM, obtained via bootstrap. **b**. Performance of amplitude cue discriminator model on Experiment 3b. **c**. Performance of spectral centroid discriminator model on Experiment 3b. **d**. Results of Experiment 3c. Experiment was identical to Experiment 3b except that the peak amplitude of each sound was set to a fixed reference, such that the two sounds on each trial were level-equated. The worse performance compared to Experiment 3b is unsurprising as the resulting stimuli are physically incorrect. **e**. Performance of amplitude cue discriminator model on Experiment 3c. **f**. Performance of spectral centroid discriminator model on Experiment 3c.

### Experiment 3c: Mass discrimination across surfaces with level-equated sounds

The results of Experiments 3a and 3b along with the observer model results indicate that human judgments are not exclusively driven by the acoustic cues that are associated with mass (amplitude and spectral centroid), at least not when humans are forced to separate mass from other object variables. To further test this idea, we ran a control experiment (Experiment 3c) in which the amplitude of the two sounds was equated, eliminating one of the cues that might plausibly be used. In this case performance was worse overall, but was still well above chance for the two conditions that were otherwise realistic (Recorded and Full model), while notably worse for the conditions from lesioned model variants (Fig. 9d). This task obviously cannot be performed by the amplitude cue discriminator (Fig. 9e), and is also not explained by the centroid discriminator (Fig. 9f), which is more accurate than humans for large mass-ratios but less accurate for low mass-ratios, and which again does not differ between conditions. We note that the relative sound levels here were physically incorrect, which in some contexts causes humans to mis-identify sounds^42^, and could explain the worse performance relative to the realistic case where there are level differences between sounds. This result nonetheless reinforces the idea that listeners are not basing judgments on amplitude cues alone.

Overall, the results indicate that listeners are able to separate the contribution of object resonances from that of the force exerted between objects, in this case as is determined by the mass of the impacting object, and that they rely on knowledge of real-world impulse response regularities to do so.

### Experiment 4: Perception of damping

A second everyday scenario in which perceptual inferences are ill-posed involves the perception of external damping^39^. External damping occurs when one object abuts another, absorbing some fraction of the other object’s vibrations. External damping is distinct from the internal damping in which energy dissipates within an object at rates that depend on the material (reflected in the material-dependent decay rates seen in our impulse response measurements, and in prior theoretical accounts of material perception^14,16,18,39^). In this paper, we use the term damping to refer to external damping.

Nearly all everyday objects are externally damped to some extent, as gravity usually causes them to rest on something, but some conditions produce much more damping than others. Consider, for instance the sound of tapping a coffee mug sitting on a table vs. held firmly in someone’s hand. Anecdotal experience suggests we are able to discern the degree of external damping to some extent, but the acoustic basis of any such abilities was unclear.

We began by measuring the acoustic effects of external damping on object resonances. We conducted two experiments: one in which we recorded impact sounds produced by tapping an object that was either resting freely on a table or being held tightly in the experimenter’s hand (Supplementary Fig. 7), and another in which the degree of damping was parametrically manipulated by clamping different fractions of a board and then measuring its impulse response (Fig. 8a&b). In both cases the primary effect of damping was to lower the decay time of the object resonances captured by the object’s impulse response.

This acoustic effect raises the possibility that it could be challenging to disentangle the effects of external damping from the effects of material, because both of them influence the decay time of object resonances, potentially making the inference of damping ill-posed. For instance, an undamped piece of wood might exhibit similar decay times to a damped piece of metal (Fig. 8c). However, with prior knowledge of the distribution of impulse responses, one might be able to use the frequencies of the resonances to estimate the expected decay rates in the absence of damping, enabling the effect of material to be factored out and thus damping to be more accurately estimated. We thus sought to assess whether humans could identify damped objects from sound, and whether the ability to do so would be dependent on the sounds being faithful to the regularity of real-world object impulse responses.

Participants heard a single impact sound and judged whether the object making the sound was resting freely on a table or held tightly in someone’s hand (Fig. 8d). We simulated different degrees of damping by decreasing the mode decay times (b_j_ in the parametric impulse response model described earlier) by different proportions. We synthesized sounds in six of the seven conditions from Experiment 3: two that obeyed real-world impulse response statistics (Recorded and Full model), and four that violated them in some way. If listeners assess damping simply via the absolute decay time of a sound, performance should be similar for all conditions, as the average RT60 of the sounds changed by the same amount, a prediction we confirmed with a cue discriminator model based on RT60 (Supplementary Fig. 8). We omitted the Inverted distribution condition that was part of Experiment 3 because it increased the RT60 differences between damped and undamped sounds (affecting the observer model’s performance), and thus was not an appropriate test of our hypothesis. Because this was a single-interval task, performance was quantified as d-prime.

As shown in Fig. 8e, participants were able to determine whether an object was externally damped, but were substantially better for the two conditions with naturalistic impulse responses. We again summarized the results with a threshold for each condition (here the amount of simulated damping needed to produce a d-prime of 1 in each condition; Fig. 8f). The four unnatural conditions all produced significantly worse performance than the natural conditions (p<=.003 for each paired comparison with the Recorded condition, via permutation test, statistically significant after Bonferroni correction; difference in mean threshold >= 0.092).

Moreover, for the conditions that altered mode statistics, thresholds varied inversely with the likelihood of the impulse responses under the real-world distribution, as in Experiment 3 (Fig. 8g). These results indicate that listeners rely on implicit knowledge of the distribution of object resonance properties to be able to discern when an object is damped. This experiment thus provides another example of human abilities to separate distinct physical causes of sound, using internalized knowledge of the world.

## Discussion

We investigated human perceptual inferences of physical variables from sound, leveraging measurements of real-world object acoustics. We conducted the first large-scale set of object impulse response measurements, revealing systematic regularities in the way that objects of different materials and sizes produce sound when struck. We then developed a physically-based generative model using these measurements. The model allowed us to sample and synthesize new instances of impact sounds that sounded realistic and that conveyed physical variables such as material and mass to human listeners, providing a tool with which to probe their perception. We validated the model with both physical measurements of impact sounds and with perceptual experiments showing that the model components were sufficient and necessary to render realistic impact sounds. We then asked whether listeners are able to separate the contributions of the many distinct physical variables that contribute to any single sound. We found that human listeners were able to perform this separation, as exhibited by the ability to discern mass and damping despite variation in material, but that they depended on implicit knowledge of physical regularities. In particular, when object impulse responses were drawn from alternative distributions, human performance degraded in a way that was predicted by the likelihood of the impulse responses under the estimated real-world distribution, indicating that listeners use knowledge of this distribution to factor out the effects of material when judging mass and damping. Notably, the ability to infer the properties of one of the objects in an event (the mass of the impacting ball, or the presence of a hand holding the struck object) was dependent on whether the properties of the other object could be reliably inferred from resonant modes.

The results suggest that the brain has internalized aspects of the physical and statistical structure by which sound is generated by objects, and performs inference that leverages this structure to overcome the intrinsic ill-posedness of the problem. The results implicate knowledge of the distribution of object resonances, but leave open the details of how this knowledge is applied. In principle, listeners might have a rich model of the physical generative process in their heads, which they invert via Bayesian inference to estimate physical variables. However, our results are equally consistent with a wide range of alternative (approximate) inference schemes provided they make use of the distribution of object resonances. The results appear to rule out the commonly proposed idea that physical variables are identified with particular acoustic cues, as such accounts do not predict the strong dependence on whether object resonances conform to the natural distribution.

### Relation to prior work

Three main areas of prior work are closely related to our project. The first is the body of research exploring human perception of impact sounds. Much of this work has focused on deriving individual cues from physical considerations and testing whether these cues can be used to make perceptual judgments^13–17,19–22,24–26,28^. We built on this work in three main respects. First, we studied the consequences of the ill-posed nature of the inference problem faced by observers in realistic settings in which multiple physical variables are unknown. This ill-posedness has been noted by others^18,23,24,39^ but has only been studied in a few settings^16,18,23,24^, typically in the context of a search for cues to one variable that might be invariant to other variables. We showed that humans are able to judge physical variables even in ill-posed conditions that require separating the contributions of the underlying objects (e.g. the dropped ball from the struck surface, or a struck object from the hand that may be holding it). By manipulating the regularities induced by physics, we showed how these regularities are key to physical judgments in such ill-posed conditions. Our results suggest that human judgments are not exclusively driven by individual cues (at least not the cues that would naturally be associated with the variables of mass and damping that we studied), and instead involve a more sophisticated inference that factors out the different variables that contribute to a sound. We also note that our results are fully consistent with the many prior results that implicated individual cues in settings in which judgements are relatively well-posed (i.e., in which only one physical variable differs between stimuli, as in Experiment 3a; Fig. 8g). Second, by building a method to synthesize arbitrarily many realistic impact sounds, we enabled experiments with realistic sounds without requiring them to be recorded by hand^43^. This innovation expands the experimental playing field by making it possible to conduct large numbers of experiments with large numbers of conditions. Third, by estimating a distribution over real-world object impulse responses, we were able to derive alternative (unnatural) distributions that deviated from it to varying extents. Having an estimate of the real-world distribution allowed us to quantify the extent of deviation in each unnatural condition (Fig. 6b), which enabled a stronger test of the hypothesis that humans make use of implicit knowledge of the real-world distribution (Fig. 8j & 10g).

A second relevant area of prior work is that of physical sound synthesis. Most contemporary techniques for synthesizing the sounds of physical events are based on finite/boundary element methods^44^. These approaches rely on precise descriptions of object geometry and material, from which numerical solutions to an object’s resonances can be derived. Such methods can produce highly realistic sounds^8–11^ when the physical descriptions they rely on are accurate, as is typically possible for simple shapes and standardized materials. However, physical descriptions (e.g. of an object’s shape or material stiffness) that are sufficiently accurate for these methods are not currently available for many everyday objects. We thus opted to instead measure impulse responses for a large set of everyday objects, and then to characterize their acoustic properties in statistical terms. We found that this statistical characterization was sufficient to synthesize realistic sounds. This approach was well-suited to our scientific purposes, in that it exposed the ill-posed nature of the underlying inference problems, revealed regularities that might be used to solve these problems, allowed us to violate these regularities, and enabled us to synthesize large numbers of realistic sounds. However, our approach gives us no way to synthesize a sound for a specific new object with high accuracy. It is thus complementary to the dominant current approach to physical event sound synthesis.

Another research area within physical sound synthesis – procedural audio synthesis for computer game simulations – has grown rapidly in recent years^45^. Such algorithms seek to synthesize realistic sounds with methods efficient enough to work in real-time. Many such algorithms use a hybrid approach similar to our generative model, in which some causal factors are simulated via physics (e.g., impact forces). These causal factors are often used to filter pre-recorded impact sounds, or pre-computed impulse responses^10^. To our knowledge, our model is the first to generate impulse responses by sampling mode statistics, and thus complements existing tools in this domain.

A third important area of prior work is the classical theory of acoustics. Acoustical theory predicts that objects will have resonant modes, that the amplitude of these modes will vary with the point of measurement and the point at which force is applied, and that energy in these modes will decay exponentially^5^. Predictions of mode properties can be derived for simple shapes^6^. However, it is less clear whether the predictions of classical theory hold for everyday objects that sometimes have complicated shapes and imprecisely specified materials. Our contribution relative to this body of prior work was to make empirical measurements from everyday objects and to show that the key predictions of classical physics largely hold for such objects (Figs. 2, 4 and 8, and Supplementary Figs. 1-5, 7, 9-10). These measurements in turn enabled us to build synthesis models to generate new examples of impact sounds that can be used for experimental purposes.

### Limitations

The survey we conducted of object acoustics was more comprehensive than previous measurements, but was nonetheless limited by practical constraints. We attempted to sample from the distribution of everyday experience by asking humans to name objects they had heard produce impact sounds, but we had no way of validating whether the human reports were accurate. It is possible that humans are more likely to name some types of objects than others for reasons unrelated to the frequency with which they encounter them making sounds, which could skew the results. Our sample sizes in each size/material category were also constrained both by what humans reported hearing, and by the objects we could obtain and measure. We have some reason to think that the sample sizes were sufficient for our purposes, in that material discrimination of sounds synthesized from the generative model (which was based on statistics obtained from the measured impulse responses) showed similar patterns of confusions to those for recorded impulse responses (Fig. 5c&7d). However, the sample sizes could be a limitation in some contexts.

Our survey allowed us to make measurements from the sorts of objects that humans actually encounter, rather than being entirely reliant on simple shapes (bars, plates etc.) that have been the focus of prior studies. One cost of this approach is that if object shapes are not independent of material, as is likely to be the case in real-world settings, a real-world object survey will inevitably confound effects of shape and material. This was clearly the case to some extent for our measurements. For instance, some material classes did not show the expected effect of object size on mode frequency (lower frequencies for larger objects; Supplementary Figs. 3-5). This almost surely reflects differences in the object shapes that people reported encountering in different size classes for particular materials. For example, small ceramic objects are likely to be mugs, whereas large ceramic objects are likely to be tiles. In addition, because of the heterogeneity of shapes that were involved, we are unable to analyze effects of shape itself. The results are plausibly reflective of the objects that people actually encounter in life, but come at the cost of obscuring the contributions of different object variables. Our results are thus complementary to traditional approaches with simple shapes that help identify the effects of particular variables but that do not consider real-world objects.

We were also limited by our measurement method. It was not possible to measure impulse responses from very small objects (because there was not enough surface area to attach the speakers and microphones to the object). We also did not attempt to comprehensively make measurements from very large objects, because our pilot experiments made it clear that our apparatus could not supply enough power to measure an impulse response that was sufficiently far above the noise floor for our purposes. This difficulty was compounded by the fact that very large objects tend to be very highly damped compared to smaller objects (evident in Supplementary Fig. 2, for instance). We also note that the measurement apparatus itself inevitably influences the measurement by imposing some degree of damping on the object being measured. The degree of damping is determined by the surface area of the microphones and speaker used for the measurement, and thus is probably not more than what would result from an object resting on a table, but we did not quantify this effect.

We also made the choice to characterize objects exclusively by their impulse responses. In particular, although an object’s stiffness also influences the sound it produces on impact, we estimated stiffness from theoretical calculations based on known material-specific constants rather than measuring stiffness in every object. This choice was partly due to practical constraints. Measuring stiffness required recording impact sounds and then fitting our parametric impact sound model to the recording (see Supplementary Fig. 10 for an example), and it was impractical to obtain clean recordings for each of the objects in the survey (this would have required transporting each of them into a sound booth, whereas the impulse response measurement could be made in the object’s home environment). We view this choice as reasonable on the grounds that the impulse response is a high-dimensional quantity that contains substantially more structure (70 parameters in our parametric model) than the single scalar value captured by the stiffness coefficient. In addition, the experiments we conducted to validate the stiffness coefficients used in our generative model indicate that these assumed coefficients are close to the actual values.

Our generative model ignores the issue of sound radiation. In real objects, different points on an object’s surface vibrate differently (reflected in the variation in impulse responses revealed in the analysis of Supplementary Fig. 1), such that for objects of sufficient size the sound heard by an observer might depend on their position relative to the object. We neglected such issues on the assumption that they contribute modestly to everyday experience, but they likely matter in some contexts, and should be studied. The generative model also assumes linearity. This is a good assumption for rigid objects, but cannot account for objects that bend or break^46^.

We have also limited our experiments to situations where only one of the two impacting objects radiates sound. This situation is an appealing starting point due to its relative simplicity, and the insights gained should be of general relevance to more complicated scenarios. Future experiments in which both objects radiate sound could address the question of whether it is possible to separate the contribution of multiple object impulse responses, for instance to infer both materials of the two impacting objects.

We assume that the knowledge of object acoustics that is evident in our experimental results is “intuitive” in the same sense as other knowledge of ecological optics or acoustics that constrains human perception. Most humans do not have explicit knowledge of the physics underlying how sounds or images are generated, such that judgments that leverage this knowledge do not usually involve conscious reasoning about the underlying principles. Rather, the knowledge is baked into perceptual processes that tend to be encapsulated and cognitively impenetrable, and that are thus evident only indirectly. However, we note that we did not test the extent to which our participants had any awareness of the basis of their judgments, and so do not have definitive evidence of the extent to which their knowledge of acoustics is fully implicit.

### Future directions

Our generative model and perceptual results raise the possibility of a model of inference that could account for human perception. Inference in generative models has been used to account for the perception of faces^47^ and everyday auditory scenes^48^, and it would be natural to extend such approaches to physical event perception using the tools developed in the present work. Being able to synthesize realistic sounds for physical events also opens the door to studying the perception of such events in richer conditions. For instance, the ability of humans to separate the physical variables underlying impact sounds raises the question of whether they can also separate these variables from other environmental effects, in particular reverberation^33,37,49,50^, spectral coloration^51^ and distance^42,52^. In addition, object interactions often involve scraping, rolling, and sequences of impact sounds. Progress in rendering such sounds^53^ should enable studies of auditory perception that extend beyond single impacts to more extended and realistic sequences.

The perception of physical events is also a promising problem with which to study cross-modal integration. “Intuitive physics” has also been studied in the context of exclusively visual sensory input^54–59^. Yet the perception of such events is typically multi-modal, with sight and sound often providing complementary sources of information^60,61^. The development of simulators that can render relatively realistic audio and video for physical object interactions^62^ should enable progress in this direction.

The general problem that we study here – the perception of physical variables from sound – seems likely to have broad relevance to other species. As such, it may be a good model problem for auditory neuroscience. We hope the tools that we have developed in this paper will help to stimulate additional research on this important aspect of everyday hearing.

## Methods

### Informed consent

All participants gave informed consent. The Massachusetts Institute of Technology’s Committee on the Use of Humans as Experimental Subjects (COUHES) granted approval for all experiments.

### Documenting the objects heard in everyday human experience

#### Surveying human experience with object sounds

We sought to measure from a representative sample of the objects people experience via contact sounds in their everyday life. We conducted two online surveys asking participants to report the impact sounds they had recently heard and to report the colliding objects involved. All experiments were conducted on Amazon’s Mechanical Turk platform.

Each survey asked 100 people to report the objects involved in the most memorable recent impact sound they could recall. The first survey focused on impact sounds heard within the last hour, while the second survey expanded the timeframe to include memorable impact sounds from the entire previous day (a 24-hour period). We hoped the two surveys together would reflect the range of different impact sounds encountered (or at least noticed and remembered) in daily life. Both the surveys asked participants to list very commonly heard impact sounds, and to separately list the associated objects within several size categories.

The text of the questions for the first survey was as follows:

“This is a survey of common sounds in everyday scenes. Consider the sounds you have heard in the last hour. Which objects have you heard being struck (or tapped, scraped, dropped, etc.) in this time? Try to specify the material(s) of each struck object as best as you can in a few words. Please list at least one object for each size category. If you don’t recall any specific impact sounds, list objects you might expect to hear struck in this time.”

The text for the second survey was the same except “the last hour” was replaced with “the last 24 hours”.

Participants were given a text box to type answers into for each of five size categories: smaller than 4in. (10cm), 4-12in. (10-30cm), 1-3ft (30-100cm), 3-10ft (1-3m), and larger than 10ft (3m). Participants could list more than one object and many did. No examples were given, so as not to bias participants. Participants choosing their own descriptive terms for objects and material.

No geographic filters were used but participants were requested to give their location. 82% (164/200) reported locations in the United States, 35 reported locations in India, 1 did not report a location. The age and sex of participants was not recorded. As no filters were used to screen participants (except a minimum age of 18) we expect the participant population to be typical of the Mechanical Turk pool when the surveys were run (January 2020).

From the responses, we tallied the number of objects of every named material in every size class. This procedure was performed manually by the authors. Material descriptions were quite consistent across respondents, with almost all mentioned objects being labelled with one (or several) of 17 material categories (summarized in Table 1). We categorized objects labelled with multiple materials as “composite” (which we designated as an 18^th^ material category). Many such composite objects were electronic devices (e.g., phones, laptops, lamps, coffeemakers, etc.) which were usually reported as a mixture of plastic, metal, and glass. A small number of responses listed very specific materials and were re-categorized into one of the broader and more frequently mentioned classes (e.g., copper, brass, etc. to “metal”; plywood, cedar wood, etc. to “wood”; etc.). A similar categorization was performed for object names, in order to determine the number of unique objects in each material and size class.

The final survey categories are summarized in Table 1. Of 2802 responses, 475 unique objects and 18 material categories were identified. The distribution of responses was very strongly skewed towards a small subset of objects and materials in each size category. Only 107 unique objects accounted for over half of all the responses, and only 8 material categories accounted for 91% of all responses.

#### Selection of objects to be measured

Our survey of recorded object impulse responses aimed to record as many of these commonly identified objects as was practical over a range of materials and size categories. Some commonly reported objects could not be measured due to difficulties attaching the measurement apparatus (e.g., coins, paper, etc.). Other objects could not be measured because the amplifier and speaker setup could not broadcast enough power into the objects for a high-quality recording (e.g., concrete or brick walls, most of the floor materials, and many of the largest objects, etc.). For this reason, we did not attempt to make measurements for the largest size category. For some material categories (i.e., “composite” and rubber) our preliminary recordings showed that many objects in these categories were very strongly damped, with very short impulse responses that had no detectable modes and little variation beyond spectral shape (i.e., they all produced short sharp “clicks”). Moreover, the “composite” material class was highly variable in its constituent materials. Thus, despite the frequent mention of composite and rubber objects, we did not analyze the acoustics of these material categories. In addition, once we assessed the survey results and considered the constraints of the behavioral experiments we sought to run, we realized that two of the materials we intended to use in experiments (stone and cardboard) each did not have representation in one size category. To facilitate our eventual experiments we augmented the survey results with two stone objects in the third size category and one cardboard object in the first category. Ultimately, we obtained recordings of 497 objects accounting for 70% of the survey responses and providing multiple samples from 71% of the 51 non-empty material-size bins. We then excluded objects for which there was not at least one measured impulse response that met the SNR inclusion criterion (see below), or whose material was not one of the seven materials that we analyzed in this paper. This yielded 410 objects in total, and a total of 1502 impulse responses that met the inclusion criterion. All analyses in the paper are restricted to these 1502 impulse responses.

### Object impulse response measurement

#### Object impulse response measurement apparatus

The measurement apparatus consisted of a contact speaker that supplied vibrations to the object, contact microphones to measure the vibrations of the object in response to the speaker, and a soundcard that interfaced with the speaker and microphones and was connected to a MacBook Pro. We used two different sizes of speakers and microphones. Larger speakers and microphones have a flatter frequency response, and were used unless the object was too small to accommodate them.

##### Speakers

We used Dayton Audio surface thruster speakers. This type of speaker is designed to be mounted on flat surfaces and to produce sound in a wide dispersion pattern. This made it well suited to supply vibrations to an object through its surface. The speakers were either 32 mm or 64 mm in diameter.

##### Microphones

We used either EOOCVT (30 mm in diameter) or UXCELL (10 mm in diameter) piezo surface microphones, also designed to be mounted to a surface. Microphones were placed at 4-8 locations on flat portions of the sides or top of each object.

An 8-channel audio interface (Behringer U-PHORIA) was used to simultaneously record the response to the input signal from the microphones. Recordings were made using Audacity.

##### Input Signal

A noise signal consisting of the concatenation of 30 3s Golay sequences was used as the input signal. Golay sequences are complementary pairs of binary sequences (consisting of +1 and -1 elements) constructed so that their aperiodic autocorrelation functions sum to zero at all non-zero time shifts. This property enables the cancellation of measurement noise, leading to improved impulse response estimation^63^.

#### Object impulse response measurement procedure

We first cleaned the exterior surfaces of the object to be measured. We then positioned the object, with the goal of stabilizing it without excessively damping its vibrational response. This was achieved with dedicated supporting structures made of rubber that are commonly used for woodworking (Rockler Work Bench Cookie Plus work grippers). The speaker and microphones were then attached to relatively flat areas of the object surface using electrical tape and plugged into the audio interface. We attached as many as 8 microphones if there was room, but sometimes had as few as 4 for smaller objects without sufficient flat surface area. The connection to the object surface was checked by a) applying a light lateral force and assessing if the speaker or microphone wiggled at all and b) making an initial recording and ensuring that there was no buzzing. Once connected, we adjusted the speaker power and recorder gain, with the goal of maximizing the signal relative to the noise floor while avoiding distortion (rattling or buzzing caused by excessive speaker power or clipping caused by excessive recorder gain). We then broadcast the input signal and simultaneously recorded all 4-8 microphone channels using Audacity.

Once a recording was complete, we cross-correlated each channel of the recording with the input sequence to get 30 snapshots of the impulse response at each speaker-mic location pair. The impulse response measurement was the average of these 30 snapshots.

#### Justification for impulse response normalization

We opted to normalize all impulse response recordings in order to be able to arbitrarily adjust the gain on the speaker and microphones used for measurement. There were two motivations for this choice. First, to make sure every recording was adequately powered, we had to change the gain on the speaker and microphones from object to object. Specifically, being able to infer the mode parameters relied on there being “enough” power in the modes relative to the noise floor. For less resonant objects it helped to increase the gain to levels that would produce distortion in more resonant objects. Second, the distance between the surface speaker and the microphones influences the absolute level of the impulse response, but was not strongly indicative of general object properties. Speaker-microphone distances were larger for larger objects (because we sought to space the microphones out as much as possible, to maximize the representativeness of the measurements) and were also determined by idiosyncrasies of the object shape (because we could only place the speaker and microphones on flat surfaces). As a result, the absolute level of our measurements was not likely to capture anything meaningful about individual objects. However, it remains possible that there could be variation in absolute level across materials that the normalization we employed would obscure. Because absolute level is an important cue to auditory events^42^, it was important to ensure that we were not discarding this information were it present. To assess whether this was the case we conducted an experiment to measure these levels in impact sounds made from different materials.

We conducted an experiment in which we recorded impact sounds from multiple objects to assess how the initial amplitude varied across materials. Because the sound amplitude is also influenced by the amplitude of the force of impact, we held this constant by fixing the incident momentum (by using a fixed pellet dropped from a fixed height above the resonating surface).

We used a Shure SM57 microphone connected to a Scarlett 6i6 audio interface to record the sounds on a MacBook Pro in a sound attenuating booth. The microphone was placed about 70 cm away from the point of impact. We chose not to use a surface microphone for this experiment because pilot experiments revealed that they exhibited distortion given the relatively high impact force used to make the recordings.

We dropped a metal pellet weighing 50 grams on different locations over an exterior surface of the object being measured. The gain on the sound interface was held constant across all recordings (set to a level that captured the maximum dynamic range across objects without introducing clipping). We estimated the initial sound amplitude to be the maximum value of the measured sound waveform. We then plotted these the means of these amplitudes for each object (computed over the 10-15 impacts we recorded per object).

We found that these amplitudes did not systematically vary with material (Supplementary Fig. 9). Each material exhibited a range of amplitudes, depending on the object, but the range of amplitudes was similar for different materials. The main factor we found to influence amplitude was surface thickness, with thicker surfaces having lower initial amplitudes than thinner surfaces. This is expected given that the first vibrational wave inside the object has to travel though more material when the surface is thicker, absorbing more energy. Moreover, thin plates and beams, being more easily flexed in one direction, can support “bending modes”^64^ which create more surface displacement than similar intensity compression waves in thicker objects. Given that our analysis and experiments focused on effects of material (and overall object size), this experiment suggests that the impulse response normalization did not discard information that was critical for our purposes.

#### Inclusion criteria for impulse response recordings

Despite our attempts to select objects and recording gains that would permit high-quality impulse response measurements, in some cases we observed that the noise level in the estimated impulse response was high enough to potentially interfere with the parameter estimation. To systematically screen for such noise-corrupted measurements, we leveraged the fact that the impulse responses always decayed over time with an approximately exponential shape until they hit the noise floor. As a result, in the log-amplitude domain, the envelope of the impulse response waveform could be fit with an “elbow function” (a piecewise linear function that concatenated a sloping line segment with a horizontal line segment). We computed the amplitude envelope as the mean amplitudes for successive 5ms segments of the signal. The noise floor was reflected by the horizontal portion of the function. We estimated the noise floor for each impulse response relative to the peak amplitude as the height of the flat portion of the function relative to the y-intercept. We then discarded all impulse response measurements that had a relative noise floor higher than -30 dB.

### Selection of objects to include in experiments

For the behavioral experiments described below, we drew impulse responses from a subset of the full set of measurements (all impulse responses from metal, glass, ceramic, stone, wood, plastic, and cardboard in each of the four size categories).

### Physics-based synthesis model for object impulse responses

#### Modal synthesis of impulse response

Building on prior work, we model object IRs as a sum of decaying sinusoidal modes and a noisy transient^7,30,65,66^:

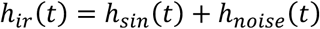

where y_ir_(t) is the IR waveform and y_sin_(t) and y_noise_(t) are the decaying sinusoidal modes and the noise transient, respectively. The key assumption is that the functional form of the decay is exponential and hence can be modeled with two parameters per mode: an initial amplitude and a decay rate. See below for description of validation of this assumption. The decaying sinusoidal modes can thus be expressed as:

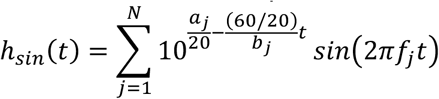

where a_j_ is the initial amplitude (in dB), b_j_ is the RT60 (in sec) and f_j_ is the frequency (in Hz) of the j^th^ mode respectively. For the purpose of this work, we only considered the 10 most salient sinusoidal modes for each IR recording (N = 10), both when analyzing and synthesizing IRs. The noise transient is modeled as a sum of decaying noise bands:

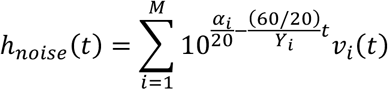

where α_i_ denotes the initial amplitude (in dB re: a maximum possible value) and γ_i_ denotes the RT60 (in sec) for the amplitude envelope on the i^th^ noise band. Unlike the sinusoidal modes, whose frequencies varied across impulse responses, the noise bands had fixed frequency limits, because the bands together spanned the entire audible spectrum. The noise bands were generated by filtering Gaussian white noise with a simulated bank of bandpass filters that were equally spaced on a frequency scale intended to match that of the human cochlea (the ERB_N_-scale^67^), with the intention of making the spectral detail equally discriminable to humans across the spectrum. Specifically, we first generated a random Gaussian noise sample and then filtered it using FIR filters whose cutoffs were equally spaced on an ERB scale. We used M = 20 noise bands, as our pilot experiments indicated that this number was sufficient to account for the perceptible variations in spectral shape in everyday IRs.

### Inferring impulse response mode parameters through analysis-by-synthesis

Because we sought to analyze distributions of impulse responses in terms of mode parameters, we first had to estimate these parameters for each measured impulse response h_m_(t). We used gradient-based optimization to estimate the parameters that best accounted for a measured impulse response (i.e., that minimized a loss function quantifying the difference between h_m_(t) and the result of the parametric function fit, h_s_(t)).

This optimization problem is challenging because many choices of loss function exhibit local minima. We settled on a multi-scale spectrogram-based loss function that quantified the difference between h_m_(t) and h_s_(t). Due to the tradeoff between temporal and spectral resolution in spectrograms, spectrograms derived from longer time windows yield better estimates of sinusoidal mode frequencies whereas those from shorter time windows are better for estimating temporal structure (here the RT60s of the modes and noise bands). We found that a multi-resolution spectrogram loss based on four different spectrograms (using time windows of 4096, 1024, 256, and 64 samples, respectively); the loss was the sum of the Huber loss^68^ between these representations of the two signals h_m_(t) and h_s_(t) [H_m,n_(f,t) and H_s,n_(f,t) respectively]. We used the Huber loss because it mitigates problems that occur for L1 and L2 losses (L1 causes slow convergence away from zero, and L2 causes slow convergence near zero). We used *δ*=3 in the Huber loss. We also added a Huber loss term between the full spectra H_m_(f) and H_s_(f) to assist the noise mode fitting:

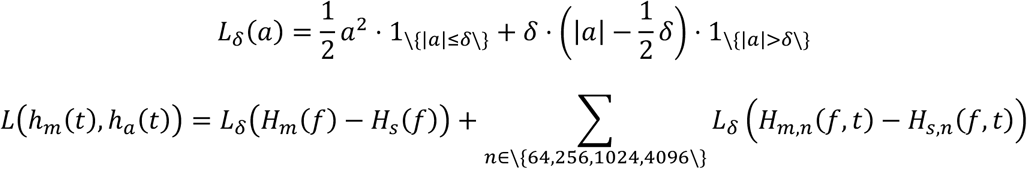

We performed gradient-based optimization in PyTorch 2.0 using differentiable audio tools contained in TorchAudio toolbox.

We found that the success of the optimization was greatly aided by “good” initial guesses for each parameter. Since the parameters have physical meaning, we used physically motivated heuristics to initialize inference.

The sinusoidal mode frequencies {f_i_} were most important to initialize well because when initialized mode frequencies are sufficiently different from their true values, small changes to the mode frequencies leave the spectrogram-based reconstruction loss unchanged, preventing gradient-based optimization from getting off the ground. To initialize the mode frequencies, we used the power spectrum of h_m_(t) to detect possible modes, using peak picking to identify the top N candidate modes with the highest average power. We applied A-weighting to the spectrum prior to peak picking to emphasize perceptually relevant regions of the spectrum. To avoid assigning multiple peaks to the same mode, we constrained the selected peaks to be at least 100 Hz apart. To avoid explaining parts of the noise transient with sinusoidal components, we required the peaks to have a prominence ≥ 2 (to reject noise bands, that were typically wider than sinusoidal modes). Peak prominence quantifies how much a peak stands out within its local neighborhood in the signal. To determine the prominence of a peak, the minimum signal value on each side of the peak is identified within the region between the peak and the closest peak of higher amplitude. The prominence is calculated as the difference between the larger of these two signal values and the peak’s height. We used the find_peaks implementation in scipy 1.14.1.

We initialized sinusoidal mode amplitudes a_i_ by uniformly sampling from [-10 dB, -30 dB]. To initialize mode RT60s b_i_ we first calculated the ‘broadband’ RT60 (the time it takes for the average power to dip by 60 dB) for the IR waveform, and then randomly initialized mode RT60s around the broadband RT60, uniformly sampling from [RT60-50ms, RT60+50 ms]. We found that this helped to ensure that the modes were fit correctly for both resonant and damped materials, and that sinusoidal modes were not erroneously accounted for by noise bands during optimization.

For the noise parameters αj, initial values were sampled from a uniform distribution according to the power (P_j_) contained in each of the cochlear frequency bands. For each band, the initial noise amplitude(αj) was initialized as a sample from U[P_j_ - 20, P_j_], with power measured in decibels. The noise RT60 γ_j_ values were initialized as samples from U[0.04 s, 0.12 s].

For gradient-based optimization during inference, we used the Adam optimizer in PyTorch 2.0. We used a learning rate of 2 × 10^-6^. Larger learning rates produced poor results, presumably because the loss landscape was not very smooth.

### Verifying discrete resonant modes of impulse responses (Figure 2d)

Although there is theoretical motivation for the assumption of discrete resonant modes^3^, it was important to test whether it held true for real-world objects. We evaluated the issue by measuring the fraction of the impulse response power that was accounted for by the N strongest sinusoidal modes. Specifically, for each record impulse response, we inferred the best-fitting modes for N = 2, 5, 10, 15, and 20 (fitting only the sinusoidal component of the parametric model, without the noise bands). We then synthesized an impulse response from these fitted modes and measured the ratio of the power of this synthesized impulse response to that in the recorded impulse response. As shown in Fig. 2d, the power accounted for by sinusoidal modes increases up to 10 modes, with little increase, on average, after that. This analysis supports the structural assumption of discrete modes that is built into the generative model.

### Verifying decay properties of impulse responses (Figure 2e)

Although there is also theoretical motivation for the assumption of exponential decaying modes^3^, it was again important to test whether it held true for real-world objects. To test the validity of the assumption, we performed the following procedure on each of the measured impulse responses. First, we calculated the power spectrum of the impulse response, and detected peaks using the same procedure described in the previous section (constraining peaks to be at least 100 Hz apart, and requiring them to have a peak prominence greater than 2). Second, we generated a spectrogram of the impulse response using with 512 fft points, window size of 128 and an overlap of 50% between consecutive windows to get sufficient resolution in time. Third, for each of the top 10 spectral peaks, we took the time-varying log-magnitude from the spectrogram at the peak’s frequency and excerpted the portion above the noise floor (estimating the noise floor as the mean log-magnitude from the last 100 ms of the impulse response, and excerpting the portion of the prior to the first point where the log-magnitude was less than or equal to this value). Fourth, we fit polynomials of degree 0, 1, 2, and 3 to the log-magnitude vs time data for each frequency using the fit function in MATLAB (which performs least-squares minimization), and calculated the goodness of fit.

As shown in Fig. 2e, this analysis showed that there was little improvement beyond degree 1, indicating that linear functions of time provided a good fit to the mode decay on a dB scale (linear functions on a dB scale are equivalent to exponential functions on a linear scale). This verified our assumption of exponential amplitude decay with time and established this as a unifying property of object impulse responses.

### Verifying broadband onset of impulse responses (Figure 2f)

We also verified that most impulse responses contain an initial broadband onset (that in our model is accounted for by decaying noise bands). For this analysis we measured the spectral flatness^69^ of successive 20-ms-long segments of each impulse response. As shown in Fig. 2f, the spectral flatness was high in the initial segment and lower for all subsequent segments, consistent with the common presence of an initial broadband onset.

### Analysis of impulse responses from the same object (Supplementary Figure 1)

To quantify the similarity between different impulse responses from the same object, we calculated the difference in the average mode parameter values for all pairs of impulse responses within an object, compared to the same quantity for pairs from different objects. We quantified the difference as the Euclidean distance between the mean mode parameters of two impulse responses. We measured the average distance for all pairs of impulse responses from the same object, for all pairs of impulse responses from different objects of the same size and material category, for all pairs of impulse responses from different objects of different sizes but the same material category, and for all pairs of impulse responses from different objects of the same size but different materials.

To quantify the extent to which different impulse responses from the same object share the same mode frequencies, we estimated mode frequencies for each impulse response and then calculated the number of pairs of mode frequencies that differed by 1% or less. We then made the same measurement for different objects from the same material category. We used this same measure to calculate the number of modes for each object that are only present in one of the impulse responses measured from the object (“orphan” modes). We then compared this number to that for groups of impulse responses of the same size as the within-object impulse responses, but with each impulse response from a different object (one across-object group matched to each object).

### Generative model of impulse responses

Our generative model of impulse responses sampled sinusoidal mode and noise parameters from distributions fitted to the estimated impulse response parameters for a particular material and size. For simplicity and ease of sampling we used Gaussian distributions that were truncated to limit the distribution to physically realizable values, and sampled modes independently. The distributions of sinusoidal modes thus captured the mean and variance of each of the three sinusoidal mode parameters (frequency, decay time, and starting amplitude) as well as the covariance between parameters. The distributions for noise captured the mean, variance and covariance of the decay time and starting amplitude of a frequency band.

#### 3D Gaussian model of sinusoidal mode parameters

After collecting all modes for a material and a size category, we removed all modes with RT60 < 0.01s and mode frequencies < 20Hz. We then computed sample means and covariance matrices, and used these as the mean and covariance for the Gaussian for that material and size category. We then truncated the density beyond the following limits: mode frequency, [20Hz, 20000Hz]; mode amplitude, [-inf dB, 0dB]; mode RT60, [0.01s, 1s].

#### 2D Gaussian models of noise band parameters

We used a similar approach to model the noise bands. The only difference was that because the noise bands in the model are fixed, we computed the means and covariances of the initial amplitude and RT60 values for each of the 20 noise bands, and had separate two-dimensional Gaussians for each band. Before estimating the means and covariances we removed noise bands with RT60<0.005s. We then truncated the densities beyond the following limits: mode amplitude, [-inf dB, 0dB]; mode RT60, [0.005s, 1s].

#### Impulse response generation

To generate an impulse response, we first selected the material and size category, then sampled 10 sinusoidal modes (each defined by a triplet of parameters) from the truncated three-dimensional Gaussian, then sampled the initial amplitude and RT60 for each noise band, and then used these values in the impulse response synthesis model described above. The sampling and distribution fitting were performed in MATLAB.

### Calculation of impulse response likelihood in different synthesis conditions

To calculate the average log-likelihood of impulse responses from a stimulus condition (described below), we used a mixture distribution obtained from a weighted sum of the distributions estimated for each material and size category, weighting each category equally. We then took each of a set of impulse responses and measured its probability under this distribution using the mode and noise band parameters, averaging the logarithm of this probability across impulse responses to obtain the results shown in Fig. 5c, 7i and 8g. For the recorded impulse responses, we estimated the mode and noise band parameters for each recorded impulse response. For synthetic impulse responses, we sampled 10,000 sets of mode and noise band parameters from each material and size category. The probability of an impulse response was obtained as the average of the mean probabilities of each mode and the mean probability of each noise band.

### Physics-based synthesis of impact sounds from impulse responses

#### Source-filter framework

We developed a source-filter model for synthesizing impact sounds from impulse responses, extending a previous state-of-the-art physical model^30^. The model assumes the collision interaction to be a linear system:

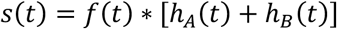

The forcing function is obtained by modeling the surfaces of colliding objects as springs, subject to physical limits on the extent of deformation. We first describe the components of the forcing function that derive from the spring model, and then describe the non-linear modification we introduced to account for physical limits on deformation.

For simplicity of analysis, assume that object A is dropped onto object B, such that object B can act as an inertial frame of reference. Under this assumption, if object A is moving with a momentum p, the resulting compression between A and B can be approximated as springs in series combination. The equivalent spring constant for such an interaction is given by

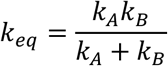

The resultant dynamics can be modeled by a half cycle of simple harmonic motion. This yields the forcing function to be 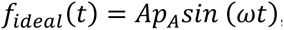, where 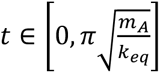, *A* is a proportionality constant, *p_A_* is the momentum of object A, and 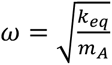.

The forcing function thus requires specifying the impact velocity and mass of the ball, and the stiffnesses of the ball and surface.

#### Stiffness

The forcing function introduced in the previous section depends on the equivalent spring constant, which is a function of the spring constants for the two constituent objects. These spring constants capture the stiffness of each object. Stiffness is a function of an object’s material, size, and shape. Because stiffness is less straightforward to measure from an object, we calculated stiffness values from known material constants and principles of solid mechanics, verifying that these values produced sounds that were very similar to those we recorded.

Stiffness is proportional to both Young’s modulus and the cross-sectional area of an object. Noting that Young’s modulus varies by orders of magnitude across the materials studied in this paper, and that the object sizes we considered also varied by orders of magnitude, we made the simplifying assumption that an object’s stiffness would be the same for all objects within a material and size category (because any variation between shapes is likely to be dwarfed by the variation across material and size). We then derived the stiffness for square plates of a size and material matched to each of the categories from the object survey.

Assuming a square plate of thickness h and side length a, with a central point load, and that is simply supported the deflection δ of the plate can be approximated by plate theory. The assumption of simple support entails that all four edges of the plate are free to rotate but not vertically deflect, with no non-zero moments.

The flexural rigidity D of the plate is:

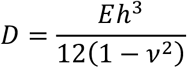

where E is Young’s modulus and *v* is Poisson’s ratio. The central deflection for a point load F on a simply supported square plate can be approximated by:

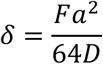

Using Hooke’s law, the equivalent spring constant k can be approximated as:

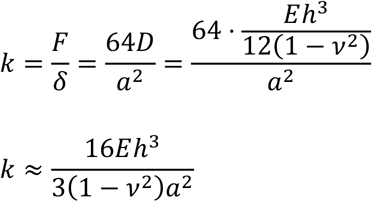

We calculated this stiffness value for side lengths of (4cm, 10cm, 30 cm and 80cm) and thicknesses of (0.5cm,1cm,1.5cm and 2cm) using material-specific values of Young’s modulus and Poisson’s ratio from engineering tables (https://www.engineeringtoolbox.com/), listed in Table 2.

**Table 2.** Physical constants used to calculate material stiffness.

| Material | Poisson's ratio | Young's Modulus (GPa) |
| --- | --- | --- |
| Glass | 0.2 | 70 |
| Ceramic | 0.25 | 215 |
| Metal | 0.2 | 135 |
| Stone | 0.22 | 42.5 |
| Wood | 0.3 | 11.5 |
| Plastic | 0.4 | 2.35 |
| Cardboard | 0.25 | 3.5 |

This calculation yielded the stiffness values given in Table 3. To calculate the equivalent stiffness needed for the impact forcing function, we took the stiffness values k_A_ and k_B_ for the materials and sizes being simulated.

**Table 3.** Stiffness constants for each material in generative model.

| Material | k (N/m) for different plate sizes (specified by side length) |  |  |  |
| --- | --- | --- | --- | --- |
|  | 4 cm | 10 cm | 30cm | 80cm |
| Glass | $3.04 \times 10^7$ | $3.89 \times 10^7$ | $1.45 \times 10^7$ | $0.48 \times 10^7$ |
| Ceramic | $11.95 \times 10^7$ | $15.29 \times 10^7$ | $5.74 \times 10^7$ | $1.92 \times 10^7$ |
| Metal | $7.03 \times 10^7$ | $9.00 \times 10^7$ | $3.38 \times 10^7$ | $1.12 \times 10^7$ |
| Stone | $2.28 \times 10^7$ | $2.92 \times 10^7$ | $1.08 \times 10^7$ | $0.40 \times 10^7$ |
| Wood | $0.69 \times 10^7$ | $0.88 \times 10^7$ | $0.34 \times 10^7$ | $0.08 \times 10^7$ |
| Plastic | $0.16 \times 10^7$ | $0.21 \times 10^7$ | $0.07 \times 10^7$ | $0.024 \times 10^7$ |
| Cardboard | $0.20 \times 10^7$ | $0.25 \times 10^7$ | $0.10 \times 10^7$ | $0.032 \times 10^7$ |

In all the experiments in this paper, we assumed the impacting object to be a small metal pellet, and so set k_B_ to the value for the smallest size category of metal. This value was the highest stiffness value across all the material-size categories. This choice had the (intentional) consequence that k_eq_ was very close to k_A_ in all cases (because k_B_ was large). If we had instead chosen the impacting object to have lower stiffness, k_B_ would have dominated the equivalent stiffness for some materials but not others, which seemed best to avoid for the sake of simplicity.

To help ensure that the assumptions underlying our stiffness calculation and the forcing function more generally were correct, we verified that the forcing function resulting from the assumed stiffness values closely approximated the sounds made by the everyday objects we sampled.

In a first test we recorded impact sounds made by pellets on objects of different materials and sizes and used the generative model to infer the stiffness of the impacted object. comparing the inferred value to the values we calculated (as described above). We used a set of four metal pellets of known masses (1.2, 5.9, 8.7, and 12 grams) and three different impacted objects from the impulse response survey (a small wood plank of size 4 x 8 x 2 cm, a large wood plank of size 20 x 71 x 2cm, and a large metal plank of size 15 x 60 x 1 cm). The motivation was both to test the assumption of linearity that yields a forcing function whose width depends on the stiffness-to-mass ratio, and to test whether the stiffness values needed to match actual impact sounds were close to the values we used in our model. We chose small metal pellets because their stiffness is high, and so should contribute negligibly to the equivalent stiffness of the impact. We recorded the sounds of 8 impacts of each pellet on each object. For each sound, we used analysis by synthesis with the generative model to estimate the equivalent stiffness (which was dominated by the impacted object due to the high stiffness of the metal pellets). We used an impulse response for each object that was measured during the survey described above (we picked the cleanest recording from the set that was recorded). We determined the stiffness value that minimized the reconstruction error given by the multi-resolution spectrogram loss defined in the above section on “Inferring impulse response mode parameters through analysis-by-synthesis”. We used a one-dimensional grid search for this purpose, first with a coarse grid sampling 500 stiffness values logarithmically spaced between 10^6^ N/m and 10^9^ N/m, and then with a finer search around the initial estimate Est_i_ in the range [Est_i_-10^0.5^, Est_i_+10^0.5^]. This analysis yielded inferred stiffness values for each object were relatively constant across different pellet masses (Supplementary Fig. 10). Moreover, the estimated stiffness values were close to the theoretical predictions that we used for stimulus generation. Both of these results help to validate the modeling assumptions.

As a second test of the overall forcing function framework, we also verified empirically that when the stiffness of the impacting object was varied, the equivalent stiffness was dominated by the softer of the two objects. Specifically, we recorded impacts made by mallets of 8 different materials when applied to a set of objects of different materials (a subset of the objects used in the impulse response survey), and then inferred the stiffness-to-mass ratio (because the mass of the mallet was not well defined, and did not matter for the purposes of the analysis because it was held constant across objects). We recorded the impact sounds using a Shure SM-50 free-field microphone. Then, using the impulse responses measured from the same objects, we inferred the object modes using the analysis-by-synthesis inference procedure described below. We then used an analogous analysis-by-synthesis inference procedure to determine the best-fitting stiffness-to-mass ratio (fixing the mode parameters). We used a 1D grid search to infer the stiffness-to-mass ratio.

This analysis confirmed that the equivalent stiffness is dominated by the softer object. Specifically, for the hardest mallets, the equivalent stiffness was determined by the object that was struck (i.e., it varied across objects). By contrast, for the softest mallet, there was little variation in equivalent stiffness across objects. This provides some additional support for the assumptions underlying the forcing function formulation that we used.

We note that for all experiments other than Experiment 1 (realism judgments), we held stiffness constant across materials to isolate the contribution of impulse response statistics. In this case (Experiments 2-5), we used a single stiffness value close to the middle of the distribution of the values in Table 3 (k_A_ = 10^7^N/m).

#### Nonlinear forcing function

We found empirically that the actual forcing function produced by everyday impacts deviated from this linear model for forces that were sufficiently large (consistent with existing knowledge in material sciences on the response of real materials to impulsive load^70–72^). We accounted for this linearity limit by using a tanh function to “soft clip” the otherwise sinusoidal impact force.

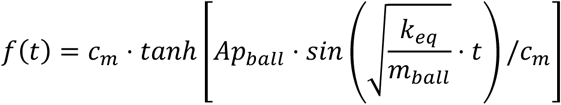

Under this model, an object is able to absorb only up to a certain amount of force that it can release in the relaxation phase. Any force above that limit is dissipated as heat and plastic deformation. The extent of the nonlinearity is governed by the incident momentum and a material-and shape-specific constant (c_m_). For instance, thicker objects will be able to act as linear springs for higher loads than thinner counterparts of the same material. We set the constant c_m_ by ear for each material, comparing synthesized sounds to impact sound recordings from the objects in the survey and finding a value that produced what to us sounded like a good perceptual match to a majority of the recordings for a material.

### Effect of damping on object impulse responses (**Fig. 8b** & Supplementary Figure 7)

Everyday objects are routinely damped via contact with other objects. We hypothesized that the effects of damping might be fairly simple to capture within our synthesis framework, which would facilitate the study of perceptual effects related to damping.

We first conducted two complementary experiments to measure the effect of damping on object impulse responses. In the first experiment, we recorded impulse responses from planks of four different materials (ceramic, glass, metal, and wood). To isolate the planks from their surroundings, they were placed on foam cones. Damping was then manipulated with weighted foam placed on both sides of the planks (Fig. 8a), varying the proportion of the plank that was encased in the foam (10%, 25%, 50%, and 75%). Impulse responses were measured with the apparatus described above, with the surface microphones and speaker placed on the exposed portion of the plank. Mode parameters were estimated for each measured impulse response and then analyzed for each material and damping condition.

As shown in Fig. 8b, the measured impulse responses revealed that the effect of damping was primarily confined to the mode decay rates: RT60 values decreased linearly as the damped area increased. By contrast, mode frequencies and amplitudes remained relatively unchanged for different damping levels.

To assess the effects of damping in a commonly occurring everyday scenario, we conducted a second experiment comparing impulse responses of objects held by a human to those of the same objects resting freely on a table. We selected comparatively resonant objects made of ceramic, glass, and wood (cups, plates and bowls). In each of the two recording conditions, we placed a surface microphone on the object at 2-3 different locations. The object was then struck using the flick of a finger, and the resulting impact sound was recorded. We then ran a mode inference algorithm on the recorded sounds, assuming a linear forcing function with a stiffness of 10^8^ N/m and a mass of 0.1kg. Finally, we analyzed the inferred mode parameters across the two damping conditions for each object.

The mode frequencies and initial amplitudes again remained relatively unchanged in this second scenario when an object is damped with a human hand, whereas the mode RT60s change substantially (Supplementary Fig. 7).

Taken together the results of these two experiments suggest that damping can be simulated by scaling the RT60s of an impulse response. For our experiments on the perception of damping, we computed the extent of RT60 variation for each material type by calculating the ratio of the clamped and unclamped RT60s from Experiment 2. We then defined a “damping ratio” to be

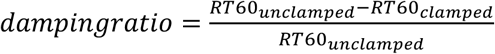

and varied this ratio from 0.1 to 0.9 to simulate different levels of damping while keeping the mode frequencies and initial amplitudes constant.

### General procedure for psychophysical experiments

To obtain large sample sizes, we ran all experiments online. Extensive research conducted in our laboratory has consistently demonstrated that online data can match the quality of data collected in traditional laboratory settings^42,73–78^ provided measures are taken to ensure standardized sound presentation, encourage participant compliance, and promote active engagement in the tasks.

We used Prolific to recruit participants. To participate, Prolific participants had to reside in Canada, the United Kingdom, or the United States, have an approval rating exceeding 95%, and be fluent in English. Target sample sizes were determined from power analyses using pilot versions of each of the experiments (see below for details). In practice, we typically slightly exceeded the target sample sizes due to the constraints of running online participants in batches.

To ensure standardized sound presentation and improve listening conditions, participants were asked to conduct the experiment in a quiet environment, minimizing external sounds. They were then instructed to adjust the computer volume to a comfortable level while listening to a calibration noise signal. The calibration was a pink noise signal peak-normalized to 0.5, and was used a reference to which stimulus levels were scaled. Each experiment then commenced with a “headphone check” to verify headphone use^79^, promoting consistent sound presentation and reducing background noise. If participants passed this check, they proceeded to the main experiment. No feedback was given in any of the main experiments.

After the main experiment, participants completed a 20-trial pitch discrimination task, the goal of which was to screen for attentiveness and task compliance. Each trial presented two harmonic complex tones whose fundamental frequencies differed by one semitone, and participants judged which tone was higher. Participants were excluded from analysis of the main experiment if their performance on the pitch discrimination task was worse than 0.85.

### Stimulus conditions for behavioral experiments

Here we describe the stimulus conditions used in the experiments. Each experiment used a subset of these conditions. All stimuli were synthesized using the impact sound model described above, and differed in the impulse responses used for synthesis (and for Experiment 3, they also differed in the mass values used in the forcing function).

#### Recorded impulse responses

Mode parameters were extracted from impulse response recordings from the survey and were used to resynthesize the impulse response. The resynthesis served to remove measurement noise, which otherwise would have differentiated stimuli generated from measured impulse responses from those from impulse responses sampled from the generative model.

#### Full generative model

The impulse response modes were sampled from the full generative model (i.e., using the fitted Gaussian distributions for a material and size category).

#### Inverted covariance

Impulse response modes were sampled from Gaussian distributions that had opposite signed off-diagonal covariance values compared to the distributions in the full generative model (that were determined by fitting to the survey data). The noise modes were sampled by reversing the assignment of the 2D Gaussians to a reversed set of ERB bands (the parameters for the first subband were assigned to the last subband, the parameters for the second subband were assigned to the second-to-last subband, and so on).

Inverted covariance with scaled variance: Impulse response modes were sampled from Gaussian distributions that had opposite signed off-diagonal covariance values compared to the distributions in the full generative model (that were determined by fitting to the survey data). In addition, the diagonal values in the covariance matrix were scaled by a factor of 5. The noise modes were sampled by reversing the assignment of the 2D Gaussians to a reversed set of ERB bands (the parameters for the first subband were assigned to the last subband, the parameters for the second subband were assigned to the second-to-last subband, and so on). The diagonal values of the individual covariance matrices for the noise modes were also scaled by a factor of 5.

#### Uniform distribution

Impulse responses modes, and noise band parameters, were sampled from uniform distributions fit to the survey data for each material and size category. The limits of the uniform distributions were set using the maximum and minimum values of the mode or noise band parameters each material and size category.

#### Inverted distribution

Impulse response modes, and noise band parameters, were sampled from distributions that were inverted about the maximum values for each mode parameter that was measured in the IR survey. This was achieved by sampling mode parameters from the Full generative model and then subtracting the parameter value from the maximum measured value for frequency, amplitude, or RT60. This shifts the mean and changes the sign of off-diagonal covariance elements.

Two modes, Gaussian: Impulse responses were synthesized from the full generative model, but using only 2 sinusoidal modes instead of 10. The noise component of the impulse response was generated as in the full generative model.

#### Linear decay

Impulse responses are synthesized from the full generative model but with linear decay in place of exponential decay. Specifically, the mode and noise band amplitude decayed linearly over time, with a linear function that dropped 60dB over the RT60 duration.

### Experiment 0: Material discrimination of recorded and synthetic impact sounds

#### Procedure

On each trial, a participant heard a single impact sound triggered by the click of a button on the trial page. Participants judged the material of the impacted object (responses selected from 7 options). The experiment consisted of 168 trials (four trials for each combination of the 3 stimulus conditions, 7 materials, and 2 size categories).

#### Stimuli

Stimuli were drawn from one of three conditions. In the first condition, stimuli consisted of recordings of impact sounds. The other two conditions were the Recorded impulse responses and Full generative model conditions described above.

The sounds for the recorded impact sound condition were recorded in a sound booth using a Shure SM57 microphone connected to a Scarlett 6i6 audio interface. The microphone was placed about 70 cm away from the point of impact. The objects were placed on foam supports (to minimize damping), which in turn were placed on a table. To generate impact sounds we dropped a metal pellet weighing 50 grams on each object, catching the pellet by hand after the first bounce. The objects were a mix of objects from the impulse response survey and substitutes that we were able to procure at the time of the recording. We drew stimuli from two size categories (10-30 and 30-100cm). The rationale for this choice was that these two sizes enabled high quality audio recordings (larger objects were often impractical to transport into our sound booth, and smaller objects often did not produce much sound with the 50-gram pellet). We recorded 10-20 impacts per object, extracted each impact, and pooled all the recorded impacts for a material and size category. We then randomly sampled from these to obtain experimental stimuli.

Each participant heard one example of the 7 material classes (metal, glass, ceramic, stone, wood, plastic, and cardboard) and 2 size categories for each condition. Stimuli for the Recorded impulse response condition were resynthesized from a recorded impulse response randomly drawn from the set of survey recordings in the desired material and size category. All sounds were synthesized using a mass of 0.05 kg, material-specific stiffness values given in Table 3, and an impact velocity of 2m/s. Each stimulus was peak-normalized to 0.5.

#### Sample size

Based on the same reasoning used for Experiment 2 (which used the same task), we targeted a sample size of 50.

#### Participants

148 participants were recruited through Prolific. Out of these, 97 participants were excluded either because they failed the headphone check, had self-reported hearing loss, did not complete the full experiment, or did not meet the performance inclusion criterion for the pitch discrimination screen. As a result, a total of 51 participants were included in the data analysis. Of these participants, 27 identified as female, 23 as male, and 1 as nonbinary. The average age of the participants was 36.7 (SD = 10.5). All participants were unique to this experiment.

#### Statistics and data analysis

To compute the confusion matrices, we pooled data across the entire participant pool for each test condition. The cell in the *i*^th^ row and *j*^th^ column of the confusion matrix was the proportion of trials in which the stimulus was generated from material *i* but was labeled as material *j*. We compared the confusion matrix for the two synthetic conditions to that for the recorded impact sound condition with a correlation coefficient.

### Experiment 1: Realism discrimination

#### Procedure

On each trial, a participant heard two impact sounds in succession, separated by 0.25 seconds. One of the sounds was synthesized using a recorded impulse response as described in the previous section, while the other was from one of six synthetic conditions. The two sounds were presented in random order. Participants judged whether the first or the second sound came from a real object. The start of each trial was triggered by the click of a button. The experiment consisted of 196 trials (7 synthesis conditions x 7 materials x 4 sizes, presented in random order).

#### Stimuli

One of the two sounds on a trial was resynthesized from a recorded impulse response that was randomly drawn from the set of survey recordings used for experiments. The other sound was synthesized from one of six conditions: Full generative model, Uniform distribution, Two modes Gaussian, Two modes uniform, Linear decay Gaussian, and Linear decay uniform. We omitted the No covariance and Inverted covariance conditions as it was clear from pilot experiments that although they altered the nature of the sampled sounds, they did not sound unrealistic (for instance, they might cause one material to sound like another, but not to sound like something that could not have occurred in the world). All of the conditions were synthesized using a mass of 0.1kg. The stiffness values were taken from the theoretical predictions given in the above section on “Stiffness”. The impact velocities were set to 2m/s to keep the forcing function in the (approximately) linear regime. Each stimulus was peak-normalized to 0.5, to help ensure that all stimuli were audible. Each participant heard each material-size combination once in each synthesis condition. There were seven materials (metal, glass, ceramic, stone, wood, plastic, and cardboard) and four size categories (all four from which we had recorded impulse responses).

#### Sample size

We sought to be well powered to detect differences between conditions using a paired t-test. Using data from a pilot experiment, we calculated the effect size for the difference between the proportion correct for the Full model and Two modes conditions, as this was a modest effect in the pilot experiment that seemed on par with the effects we would hope to detect in the main experiment. The pilot experiment was similar to the main experiment but had a different set of conditions that partially overlapped with the main experiment. The smallest effect of interest in the pilot experiment had a difference in means of 0.084, with a standard deviation of the difference between conditions of 0.24. A sample size of 67 was sufficient to achieve power of 0.8 with a significance threshold of 0.05.

#### Participants

287 participants were recruited through Prolific. Out of these, 189 participants were excluded either because they failed the headphone check, had self-reported hearing loss, did not complete the full experiment, or did not meet the performance inclusion criterion for the pitch discrimination screen. As a result, a total of 87 participants were included in the data analysis. Of these participants, 45 identified as female, 40 as male, and 1 as nonbinary. The average age of the participants was 37.4 (SD = 11.9). All participants were unique to this experiment.

#### Statistics and data analysis

We calculated the performance of the participants as a fraction of the trials they were correct on, per condition. Error bars plot the standard error of the mean across participants.

### Experiment 2: Material discrimination

#### Procedure

On each trial, a participant heard a single impact sound triggered by the click of a button on the trial page. Participants judged which of 7 materials the impacted object was made of. The experiment consisted of 224 trials (one trial for each combination of the 8 stimulus conditions, 7 materials, and 4 size categories).

#### Stimuli

Stimuli were drawn from each of the eight synthesis conditions described above (Recorded impulse responses, Full generative model, Inverted covariances, Uniform distribution, Two modes Gaussian, Two modes uniform, Linear decay Gaussian, and Linear decay uniform). Each participant heard one example of the 7 material classes (metal, glass, ceramic, stone, wood, plastic, and cardboard) and 4 size categories for each condition. Stimuli for the Recorded impulse response condition were resynthesized from a recorded impulse response randomly drawn from the set of survey recordings in the desired material and size category. All sounds were synthesized using a mass of 0.5 kg, a stiffness of 10^7 N/m, and an impact velocity of 2m/s. Each stimulus was peak-normalized to 0.5.

#### Sample size

We sought to be well powered to detect a difference between the correlations for two conditions using a permutation test. Using pilot data, we computed power empirically as a function of sample size. The pilot experiment was similar to the main experiment but had a different (partially overlapping) set of conditions. We measured the power to detect the smallest difference in correlations that was of interest (the difference between the correlation between the Recorded and Full generative model conditions and the correlation between the Recorded and Inverted covariance condition) by drawing bootstrap samples of participants of different sizes, running a permutation test on each bootstrap sample, and then plotting the proportion of significant tests (p<.05) vs. sample size. This procedure indicated that a sample size of 50 would be sufficient to yield a power of 0.8.

#### Participants

282 participants were recruited through Prolific. Out of these, 225 participants were excluded either because they failed the headphone check, had self-reported hearing loss, did not complete the full experiment, or did not meet the performance inclusion criterion for the pitch discrimination screen. As a result, a total of 57 participants were included in the data analysis. Of these participants, 30 identified as female, 26 as male, and 0 as nonbinary. The average age of the participants was 38.3 (SD = 12.1). All participants were unique to this experiment.

#### Statistics and data analysis

To compute the confusion matrices, we pooled data across the entire participant pool for each test condition. The cell in the *i*^th^ row and *j*^th^ column of the confusion matrix was the proportion of trials in which the stimulus was generated from material *i* but was labeled as material *j*. We then compared the confusion matrix for each synthetic condition to the resynthesized impulse response condition (as the latter was our best estimate of the ground truth benchmark for human material discrimination performance). We quantified the similarity between confusion matrices with a correlation coefficient. The error bars on these correlations (Fig. 6c) plot the standard error of the mean, derived via bootstrap (10,000 samples). We evaluated the statistical significance of differences in correlations using permutation tests (i.e., evaluating the probability of the observed difference under the distribution of differences in correlations measured when condition labels were permuted within participants for the pair of conditions being compared).

### Experiments 3a, 3b, & 3c: Mass discrimination

#### Procedure

On each trial, a participant heard two impact sounds in succession, separated by 0.25 seconds. Participants were told that the sounds were produced by one object dropped on a surface, and judged whether the first or the second sound came from a heavier dropped object. The start of each trial was triggered by the click of a button. The experiment consisted of 140 trials. For Experiment 3a, the 140 trials consisted of 7 synthesis conditions x 5 mass ratios x 2 material classes (hard and soft) x 2 repetitions. For Experiments 3b and 3c, the 140 trials consisted of 7 synthesis conditions x 5 mass ratios x 4 material combinations.

#### Stimuli

Stimuli were drawn from seven of the synthesis conditions described above (Recorded impulse responses, Full generative model, Inverted covariances, Inverted covariance with scaled variance, Uniform distribution, Inverted distribution, and Linear decay). We omitted the Two modes condition because it seemed plausible that the smaller number of modes would produce impair performance under any hypothesis. These seven stimulus conditions were fully crossed with five mass ratio conditions, with ratios of 1.8, 3.16, 10, 31.6, and 100. The two sounds on a trial were each generated under the same stimulus condition, with the specified mass ratio, and with the smaller of the two masses being 100 grams. All sounds were synthesized using a stiffness of 10^7 N/m and an impact velocity of 2m/s. For Experiments 3a and 3b, the sounds for the heaviest masses were peak-normalized to 0.99 (i.e., about 6 dB above the calibration noise), and all other stimuli were scaled relative to this value.

For Experiment 3a, each trial was assigned to one of two material classes: hard (metal, glass, ceramic and stone) and soft (wood and plastic). The impulse response used for the two sounds on a trial was randomly assigned to one of the materials in the class, and then the impulse response was sampled from the appropriate generative distribution.

For Experiment 3b, each trial was assigned to one of four material-combination conditions that dictated the materials from which the surfaces were sampled. The conditions drew from two groups of materials: hard (metal, glass, ceramic and stone) and soft (wood and plastic). The first condition sampled the surface materials for both sounds of a trial from the hard category. The second condition sampled both surface materials from the soft category. In both cases the two surface materials on a trial were constrained to be different, and the heavier and lighter objects were randomly assigned to the two materials. The third and fourth conditions sampled one surface material from the hard group and one from the soft group. In the third condition, the heavier object was assigned to the hard surface, and in the fourth condition, the heavier object was assigned to the soft surface.

Stimuli for Experiment 3c were identical to Experiment 3b except that the peak amplitude of each sound was set to 0.5, such that the two sounds on each trial were level-equated.

For all three experiments, all sounds were synthesized for the third size category (30-100cm), as this was appropriate for the moderately sized surface on which the objects were supposed to be dropped. Stimuli for the Recorded impulse response condition were resynthesized from a recorded impulse response randomly drawn from the set of survey recordings for the desired material and size category.

#### Sample size

We determined a target sample size for Experiment 3b, as this was the most scientifically critical of the three experiments. We sought to be well powered to detect a difference in the thresholds for two conditions using a permutation test. Using pilot data, we computed power empirically as a function of sample size using a procedure like that for Experiment 2. The pilot experiment was similar to the main experiment but had a different (partially overlapping) set of conditions. We measured the power to detect the smallest difference in thresholds that was of interest (the difference between thresholds for the Recorded and Inverted Covariance conditions) by drawing bootstrap samples of participants of different sizes, running a permutation test on each bootstrap sample, and then plotting the proportion of significant tests (p<.05) vs. sample size. By extrapolating to larger sample sizes, we estimated that a sample size of 100 would be sufficient to yield a power of 0.8. We targeted the same sample size for Experiment 3a, and targeted a somewhat smaller sample size for Experiment 3c once it became clear that the effects in Experiment 3b were fairly large.

#### Participants

Experiment 3a. 315 participants were recruited through Prolific. Out of these, 214 participants were excluded either because they failed the headphone check, had self-reported hearing loss, did not complete the full experiment, or did not meet the performance inclusion criterion for the pitch discrimination screen. As a result, a total of 101 participants were included in the data analysis. Of these participants, 44 identified as female, 53 as male, 2 as nonbinary, and 2 declined to answer. The average age of the participants was 35.1 (SD = 10.4). All participants were unique to this experiment.

Experiment 3b. 393 participants were recruited through Prolific. Out of these, 275 participants were excluded either because they failed the headphone check, had self-reported hearing loss, did not complete the full experiment, or did not meet the performance inclusion criterion for the pitch discrimination screen. As a result, a total of 118 participants were included in the data analysis. Of these participants, 52 identified as female, 62 as male, 3 as nonbinary, and 1 declined to answer. The average age of the participants was 36.2 (SD = 11.4). All participants were unique to this experiment.

Experiment 3c. 208 participants were recruited through Prolific. Out of these, 148 participants were excluded either because they failed the headphone check, had self-reported hearing loss, did not complete the full experiment, or did not meet the performance inclusion criterion for the pitch discrimination screen. As a result, a total of 60 participants were included in the data analysis. Of these participants, 30 identified as female, 27 as male, 1 as nonbinary, and 2 declined to answer. The average age of the participants was 38.7 (SD = 12.4). All participants were unique to this experiment.

#### Statistics and data analysis

We computed proportion correct for each mass ratio and synthesis condition for each participant (pooling across materials). We plotted the results as psychometric functions for each stimulus condition. The error bars plot standard error of the mean across participants. To summarize the performance in each synthesis condition, we estimated thresholds from the psychometric functions as the mass ratio producing 80% correct. We estimated thresholds by fitting a piecewise cubic spline to the psychometric function for each condition and calculating the mass ratio at which the proportion correct is 80%. For the error bars, we computed the SEM from bootstrapped samples of the participant pool. We evaluated the statistical significance of differences in the thresholds for two conditions using a permutation test (i.e., evaluating the probability of the observed difference under the distribution of threshold differences measured when condition labels were permuted within participants for the pair of conditions being compared).

#### Cue discriminator models

We tested two cue discriminator models on this experiment. The first model made the mass discrimination judgment by comparing the spectral centroid of the two sounds on each trial. The spectral centroid was computed by evaluating the mean of all frequencies in the signal weighted by the linear sound amplitude at that frequency. The second model computed an estimate of amplitude using an integration window of 40 ms (applied to the first 40 ms segment of each sound, to approximate a window of integration applied to an impulsive sound). The models compared the centroid or amplitude values for the two sounds, assigning a higher mass to the sound with a lower centroid or higher amplitude. Prior to this comparison, internal noise was added to the centroid or amplitude values, with the noise level set so as to approximately match the overall performance of the human participants on Experiment 3a (the same noise level was used for all experimental conditions).

### Experiment 4: Perception of damping

#### Procedure

On each trial, a participant listened to a single impact sound, triggered by the click of a button. Participants judged which of the following two scenarios best explains the condition of the struck object: sitting on a table, free to vibrate; or held by a person with their hands. The experiment consisted of 300 trials (6 synthesis conditions x 5 damping conditions x 5 materials x 2 object sizes).

#### Stimuli

Stimuli were drawn from six conditions (Recorded impulse responses, Full generative model, Inverted covariances, Inverted covariance with scaled variance, Uniform distribution, and Linear decay). We omitted the Two modes condition because it seemed plausible that the smaller number of modes would produce impair performance under any hypothesis. The Inverted distribution condition was omitted because we found the observer model based on RT60 performed better in this condition compared to the others, which made it non-diagnostic of our hypothesis. These six stimulus conditions were fully crossed with five damping conditions. The five damping conditions were generated by scaling the mean RT60 values of the marginal distributions for mode RT60s by different proportions: 1 (undamped), 0.75, 0.5, 0.25, or 0.1. The damping ratios plotted in Fig. 8e&f is one minus this proportion. Stimuli were drawn from five materials (glass, ceramic, metal stone, and wood) and the two smallest size categories. Each material and size was used once in each combination of synthesis and damping condition. Stimuli for the Recorded impulse response condition were resynthesized from a recorded impulse response randomly drawn from the set survey recordings from the specified material and size category. All sounds were synthesized using a mass of 0.5 kg, a stiffness of 10^7 N/m, and an impact velocity of 2m/s. Each stimulus was peak-normalized to 0.5.

#### Sample size

We sought to be well powered to detect a difference in the thresholds for two conditions using a permutation test. The procedure was identical to that for Experiment 3. This procedure indicated that a sample size of 50 would be sufficient to yield a power of 0.8.

#### Participants

153 participants were recruited through Prolific. Out of these, 98 participants were excluded either because they failed the headphone check, had self-reported hearing loss, did not complete the full experiment, or did not meet the performance inclusion criterion for the pitch discrimination screen. As a result, a total of 55 participants were included in the data analysis. Of these participants, 30 identified as female, 22 as male, and 3 as nonbinary. The average age of the participants was 37.4 (SD = 11.9). All participants were unique to this experiment.

#### Statistics and data analysis

We pooled the responses for each participant across materials. We then computed a false alarm rate for each synthesis condition from the undamped condition, and a hit rate for each combination of synthesis and non-zero damping ratio. We set values of 0 or 1 to 1/24 and 23/24, respectively, and then computed d’ for each participant. We plotted the results as psychometric functions for each synthesis condition. The error bars plot standard error of the mean across participants. To summarize the performance in each synthesis condition, we estimated thresholds from the psychometric functions. We estimated thresholds by fitting a piecewise cubic spline to the psychometric function for each condition and calculating the damping ratio at which d-prime was equal to 1. For the error bars, we computed the SEM from bootstrapped samples of the participant pool. We evaluated the statistical significance of differences in the thresholds for two conditions using a permutation test, as in Experiment 3.

#### Cue discriminator model

To avoid issues associated with having to pick a criterion, we ran the model on a 2I-AFC version of the experiment. The model compared the average mode RT60 of a damped and undamped sound, and picked the sound with a shorter decay time as the damped stimulus. The average mode RT60 was computed from the decay times used to synthesize the impulse response.

## Acknowledgments

Work supported by NSF grant BCS-1921501, a McDonnell Scholar Award to J.H.M., NIH fellowship F32DC013703 to J.A.T., and a K. Lisa Yang ICoN graduate fellowship to V.A. The authors thank the McDermott Lab, Nevill Hogan, and Nancy Kanwisher for comments on an earlier draft of the paper.

## Author Contributions

J.A.T. and J.H.M. designed the impulse response measurement survey. J.A.T. conducted the human survey of sound-making objects. J.A.T. and J.S. refined the impulse response measurement method and made most of the measurements. J.A.T. designed the impact sound generative model. V.A. refined the generative model by introducing the nonlinear forcing function. V.A. conceived and implemented the gradient-based procedure for estimating the parameters of recorded impulse responses. V.A. conducted the experiments validating the assumptions in the generative model about the effects of stiffness and mass. J.A.T. conducted pilot analyses of the impulse response recordings. V.A. conducted the final version of the analysis of impulse response recordings. V.A. and J.H.M. designed and piloted Experiment 0 and made the impact recordings. J.A.T., V.A. and J.H.M. designed Experiments 1-3. J.A.T. conducted pilot versions of Experiments 1-3. V.A. conducted the final versions of Experiments 0-3, and the control experiments for Experiment 3. V.A. and J.H.M. designed Experiment 4 and the experiments measuring the acoustic effects of damping. V.A. conducted Experiment 4 and the experiments measuring the acoustic effects of damping. V.A. analyzed all behavioral experiments. J.H.M. and V.A. drafted the initial version of the manuscript, with input from J.A.T. V.A. made the figures, with guidance from J.H.M. V.A., J.A.T., and J.H.M. edited the manuscript.

## SUPPLEMENTARY INFORMATION

**Figure 10.**
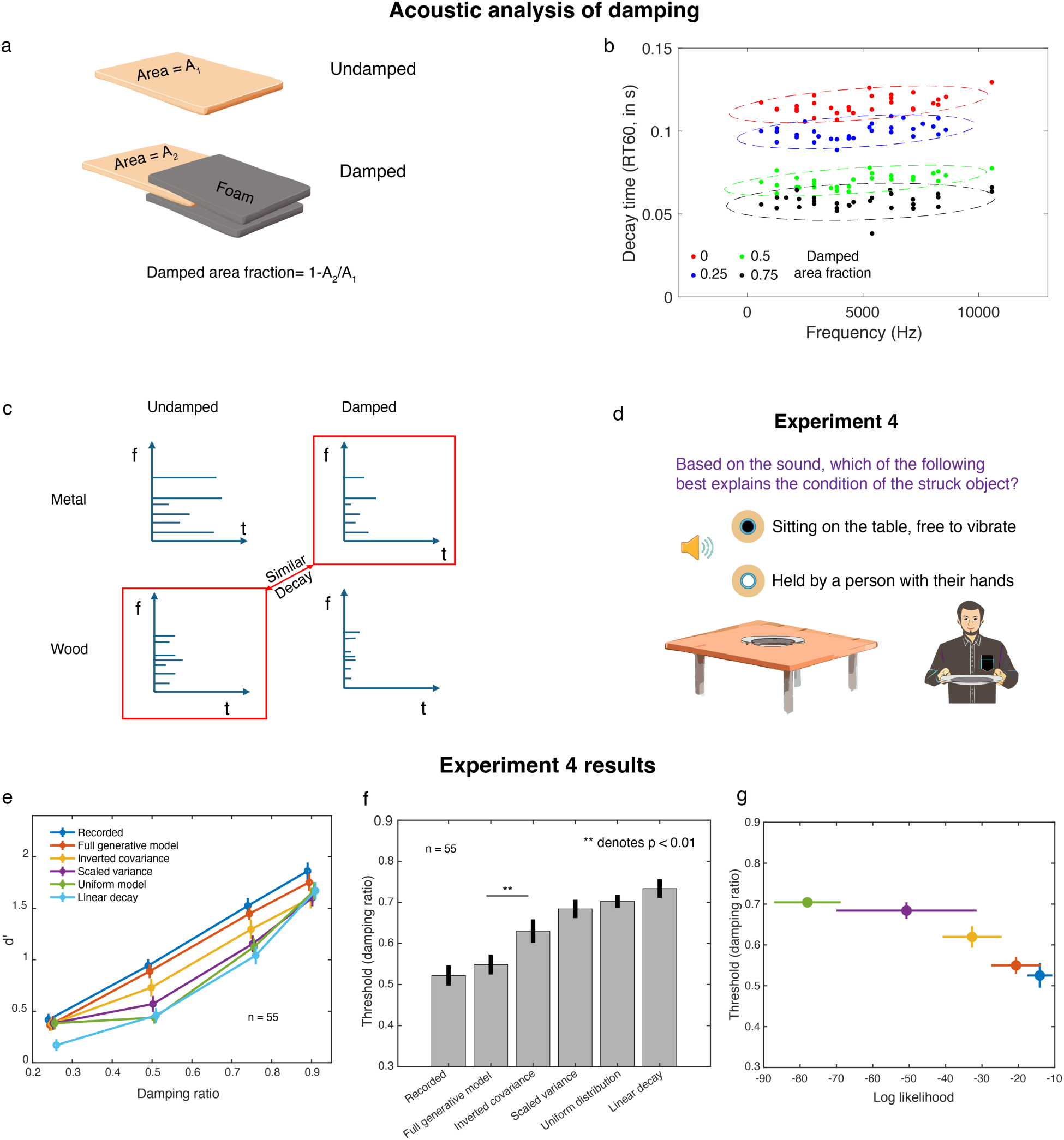
Perception of damping. **a**. Experiment measuring acoustic effects of parametrically manipulated damping. We measured impulse responses from a board that was damped to varying extents by surrounding foam, varying the amount of the board that was clamped in foam. **b**. Acoustic effects of parametrically manipulated damping. We estimated the frequencies and decay times of the impulse response modes in each of four damping conditions. Colors correspond to the fraction of the board that was damped (see panel a). Decay time decreases gradually with the extent of damping, but mode frequencies are largely unaffected. Dots plot parameter estimates for individual modes. Ellipses plot 95% confidence contours to Gaussian distributions fit to the mode parameters. **c**. Rationale for experiment: the decay times of a damped object of a resonant material (e.g. metal) might be similar to the decay times of an undamped object of a less resonant material (e.g. wood). This ill-posedness could be overcome using knowledge of the material-driven dependence of mode decay and frequency. **d**. Task of Experiment 4. Listeners heard a single impact sound and judged whether the impacted object was resting freely on a table vs. being held firmly in someone’s hand. The experiment manipulated the simulated degree of damping and the extent to which the underlying impulse responses were faithful to the physics-induced regularities of real-world impacts. **e**. Damping discrimination as a function of the amount of simulated damping, for human participants tested on the six different stimulus conditions. Error bars plot SEM. **f**. Discrimination thresholds in each condition, defined as the damping ratio resulting in d-prime of 1. Error bars plot SEM. **g**. Comparison of discrimination thresholds from Experiment 4 with log-likelihood of impulse responses. The linear decay condition is omitted as the impulse response parameters were taken from the Full model, and thus do not reflect the deviant features of that model variant. Error bars on thresholds plot SEM, derived via bootstrap. Error bars on log likelihood plot SD across impulse responses.

**Supplementary Figure 1.**
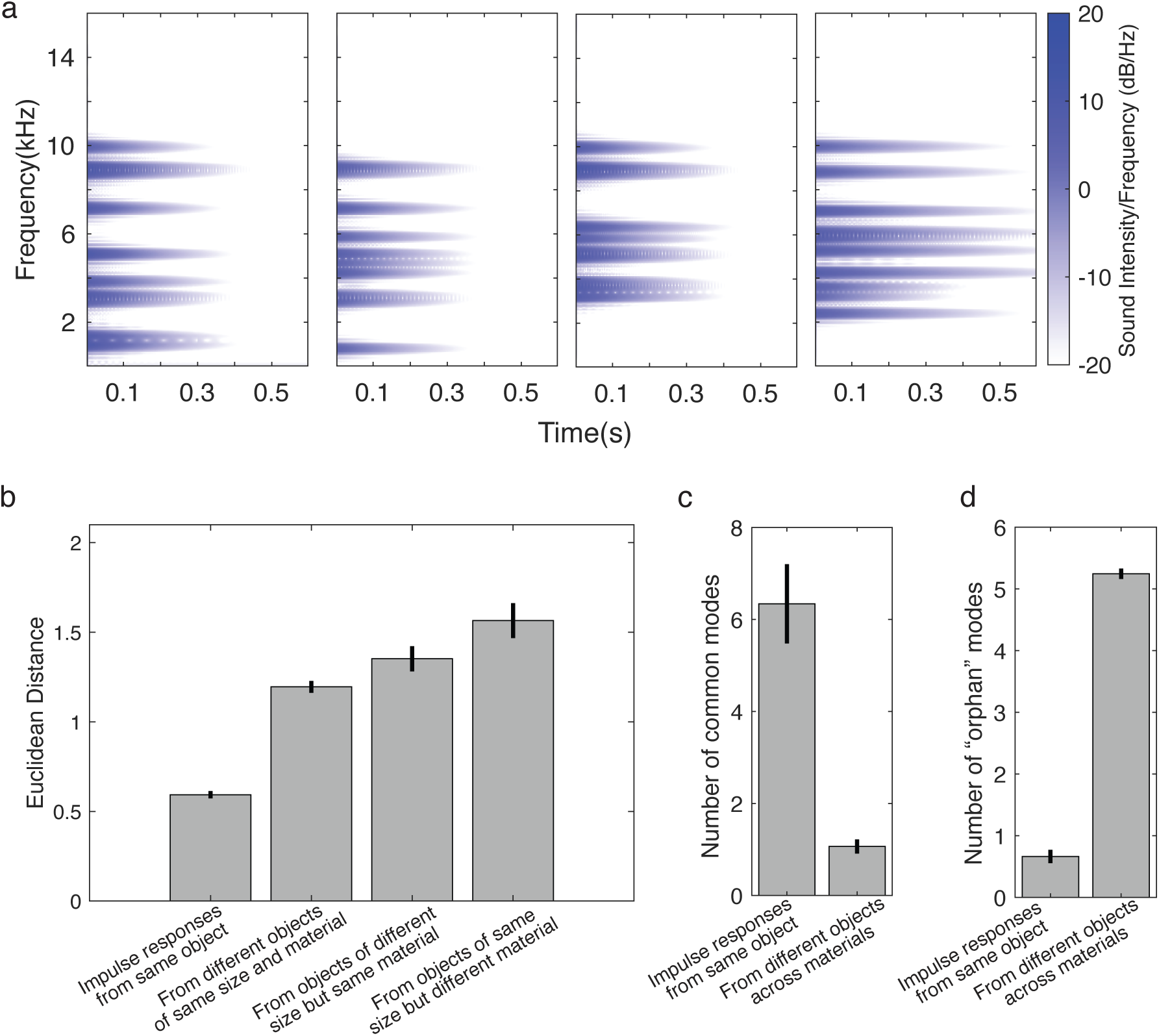
Comparison of different impulse responses from the same object. a. Spectrograms of different impulse responses measured from the same object. Visual inspection suggests that the different impulse responses share mode frequencies. b. Difference in the average mode parameter values for all pairs of impulse responses within an object, compared to the same quantity for pairs from different objects. We quantified the difference as the Euclidean distance between the mean mode parameters of two impulse responses. Graph plots the average of this distance for all pairs of impulse responses from the same object, for all pairs of impulse responses from different objects of the same size and material, for all pairs of impulse responses from different objects of different sizes but the same material, and for all pairs of impulse responses from different objects of the same size but different materials. c. Analysis of the extent to which different impulse responses from the same object share the same mode frequencies. We estimated mode frequencies for each impulse response and then calculated the number of pairs of mode frequencies that differed by 1% or less. By this measure, pairs of impulse responses from the same object tend to share a considerable number of modes (6.3 on average, SD = 0.91), whereas impulse responses from different objects do not (1.04 on average, SD = 0.14). We used this same measure to calculate the number of modes for each object that are only present in one of the impulse responses measured from the object. There were very few such modes (0.66 on average, SD = 0.11). By comparison, “orphan” modes were relatively common from groups of impulse responses drawn from different objects (5.14 on average, SD = 0.080).

**Supplementary Figure 2.**
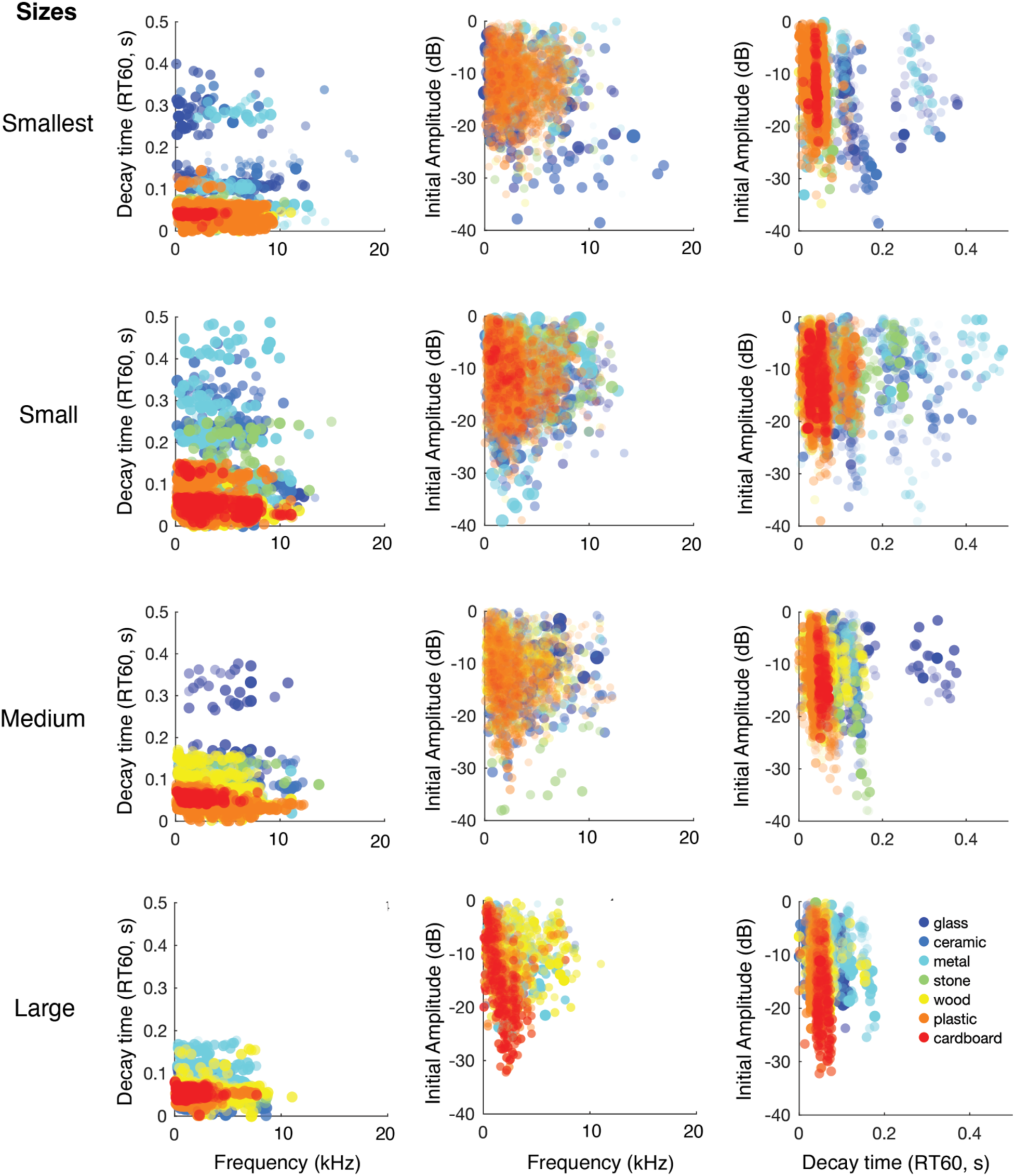
Mode parameters for all material and size categories. Each row presents the objects in one size category. Saturation of dots represents the value of the unplotted mode parameter (for instance, on the panels plotting decay time vs. frequency, the dot saturation represents initial amplitude).

**Supplementary Figure 3.**
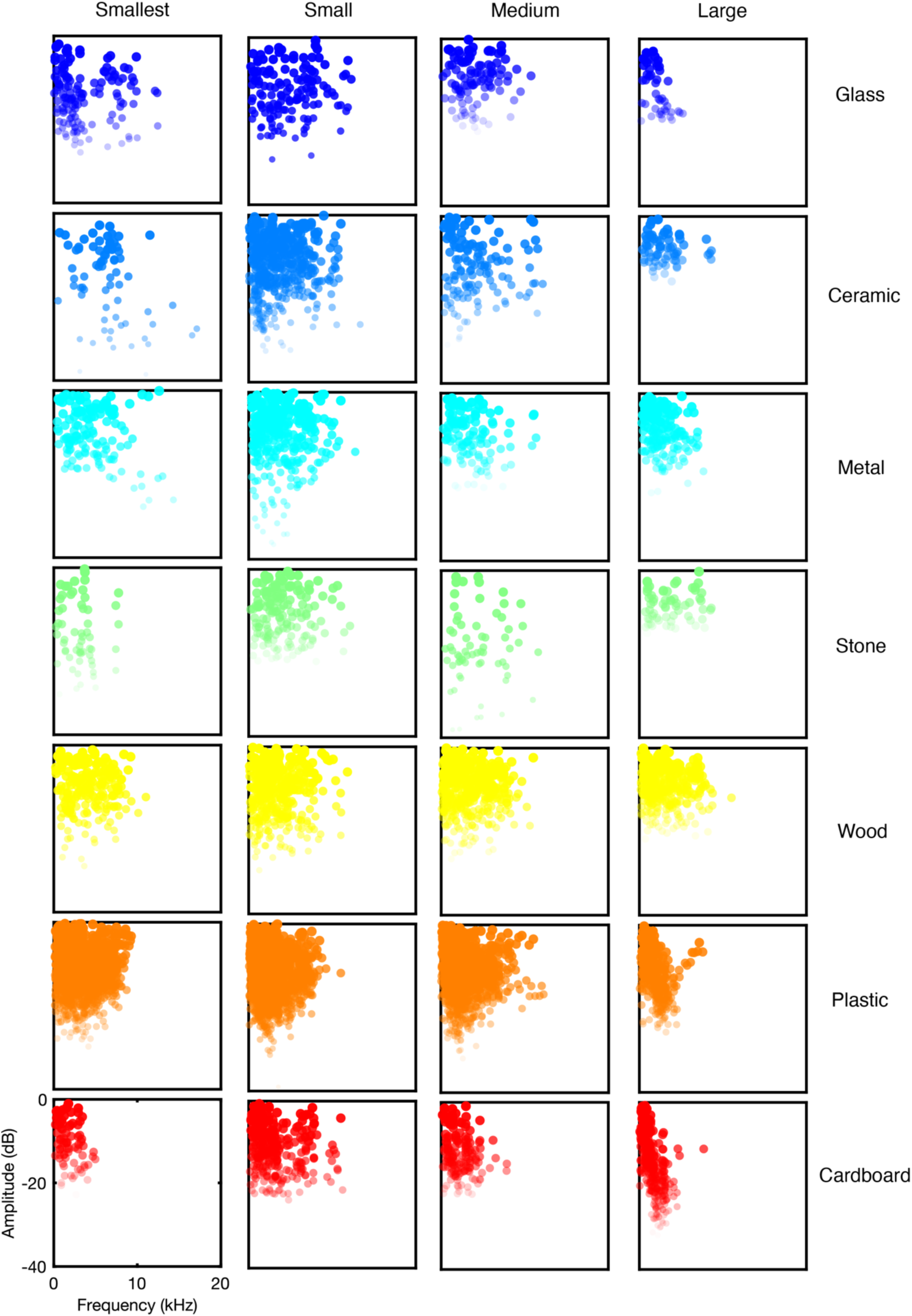
Mode amplitude and frequency for each material and size category. Same conventions as Supplementary Figure 2.

**Supplementary Figure 4.**
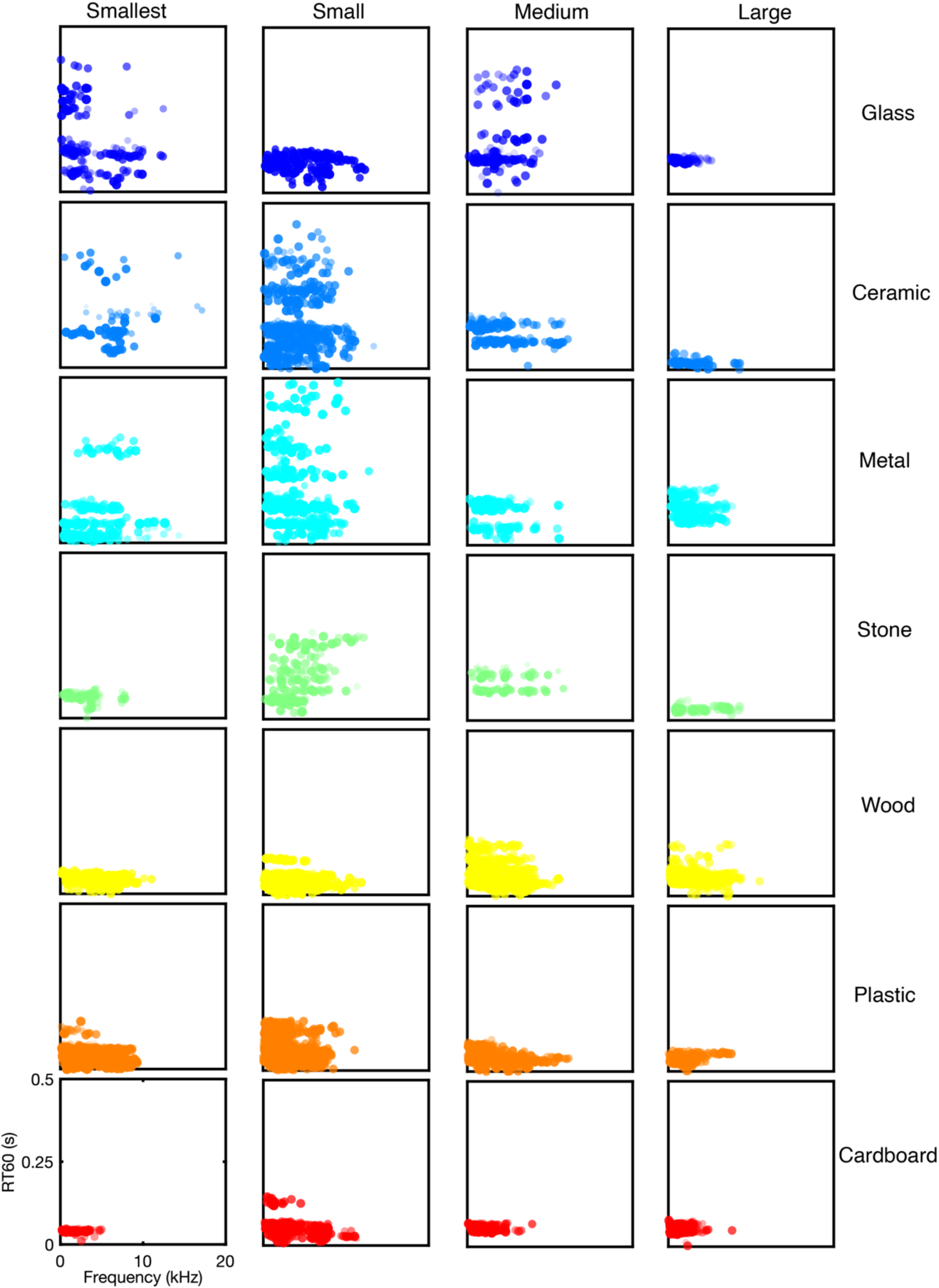
Mode decay time and frequency for each material and size category. Same conventions as Supplementary Figure 2.

**Supplementary Figure 5.**
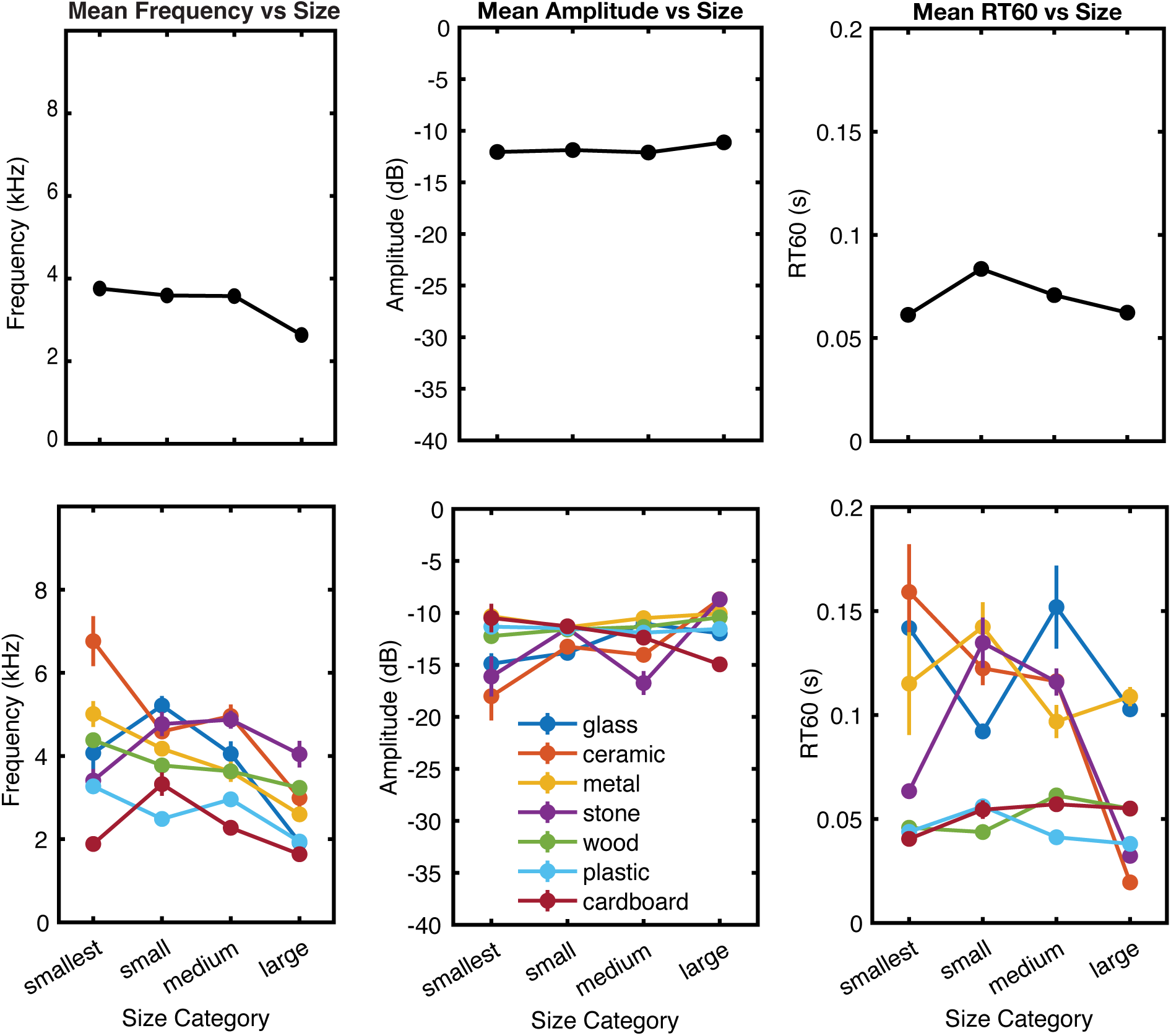
Effect of size on average mode parameters. Top shows averages across materials. Bottom shows results for individual material categories. We note that the effect of size on mode properties is in many cases confounded with object shape and damping, because different shapes and damping conditions are most common in different sizes (in the objects that humans report hearing on a daily basis). For instance, small ceramic objects are likely to be mugs, whereas large ceramic objects are likely to be tiles.

**Supplementary Figure 6.**
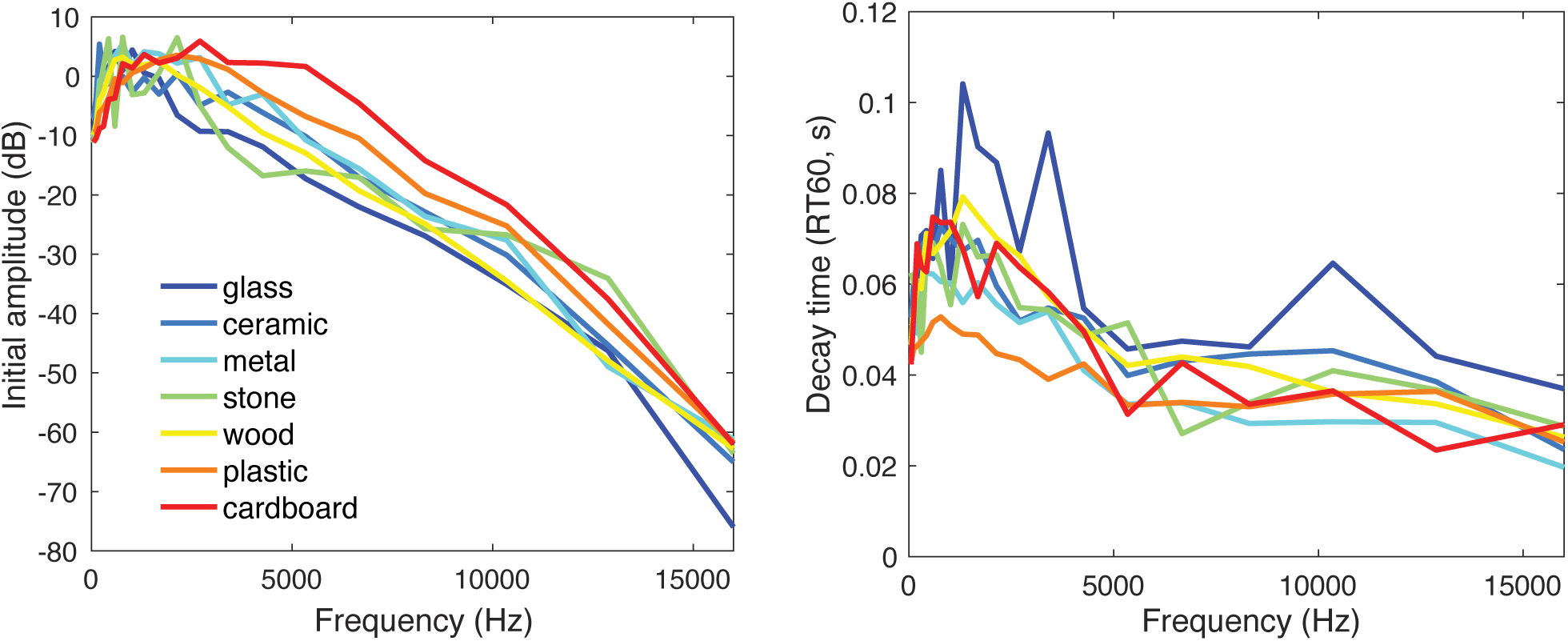
Generative parameters for noise bands of impulse responses. Note that the initial amplitudes of noise bands are higher for the two softest materials (plastic and cardboard).

**Supplementary Figure 7.**
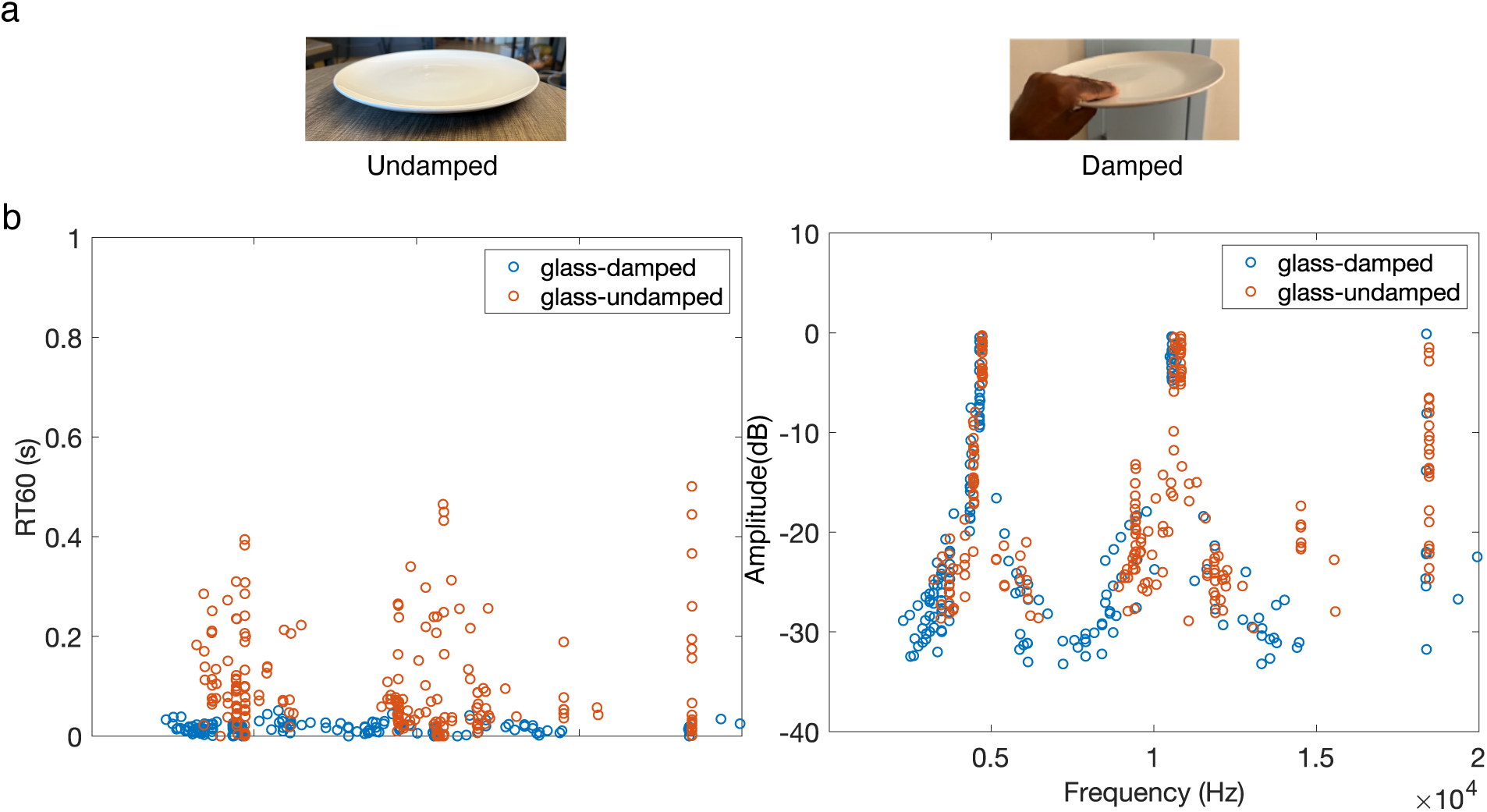
Acoustic effects of everyday object damping. We recorded impacts produced by tapping a plate that either rested freely on a table or was held by a human hand. Graphs plot the decay time, amplitude and frequency of modes measured from the recorded impacts. Decay times are much shorter for the damped plates. Frequencies and amplitudes, by contrast, remain similar.

**Supplementary Figure 8.**
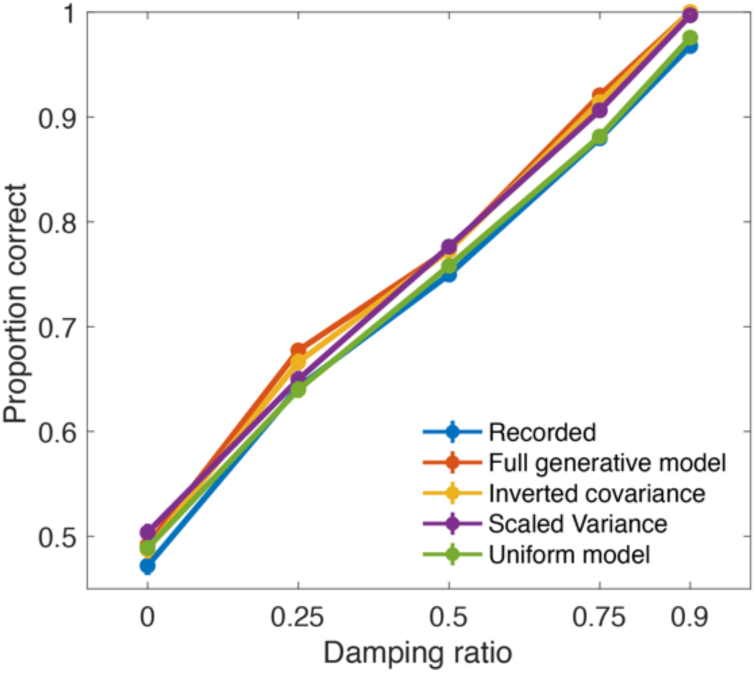
Observer model performance for damping discrimination (Experiment 4). Graph plots performance of observer model that selected the stimulus with lower decay time (average RT60 across modes) as being damped. Error bars plot SEM across 200 simulated experiment runs, and are smaller than the dot symbols. The purpose of the model was to verify that the experimental conditions did not differ in the cue that might be supposed to support performance on the experimental task. It is not intended as a model of human perception.

**Supplementary Figure 9.**
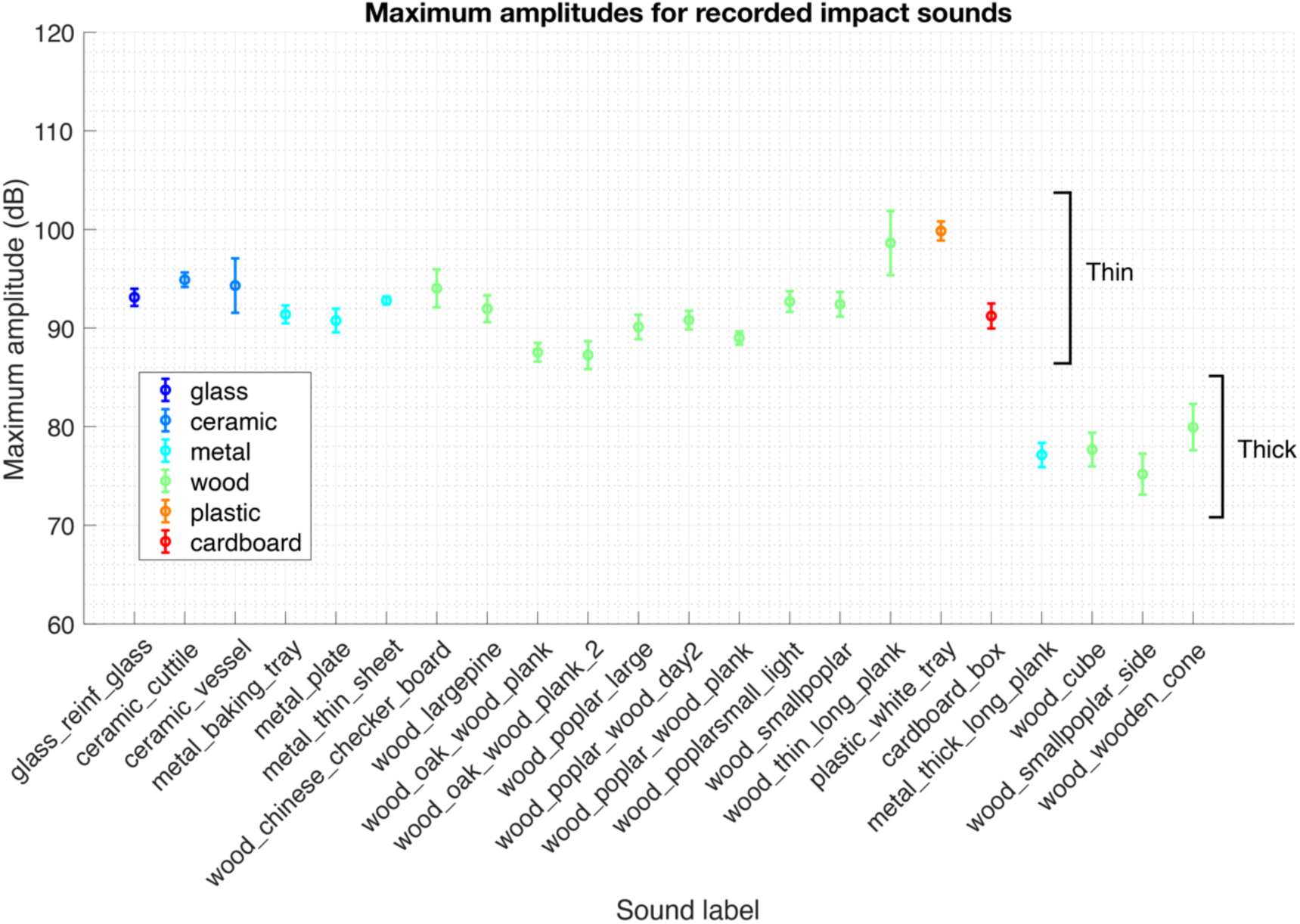
Mode amplitudes are similar across different materials. Because the impulse response measurements did not preserve absolute levels, we conducted an experiment to assess whether level varied across materials. We recorded impact sounds from objects and measured the initial amplitude of each sound. Because the sound amplitude is also influenced by the amplitude of the force of impact, we held this constant by fixing the incident momentum (by using a fixed pellet dropped from a fixed height above the resonating surface). Sounds were recorded with an external microphone, with the gain on the sound interface held constant across recordings. We estimated the initial sound amplitude to be the maximum value of the measured sound waveform. The graph plots the mean amplitude for each object computed over the 10-15 impacts we recorded per object. Error bars plot SD. Amplitudes did not systematically vary with material. The main factor we found to influence amplitude was surface thickness, with thicker surfaces having lower initial amplitudes than thinner surfaces. Given that our analysis and experiments focused on effects of material (and overall object size), this experiment suggests that the impulse response normalization did not discard information that was critical for our purposes.

**Supplementary Figure 10.**
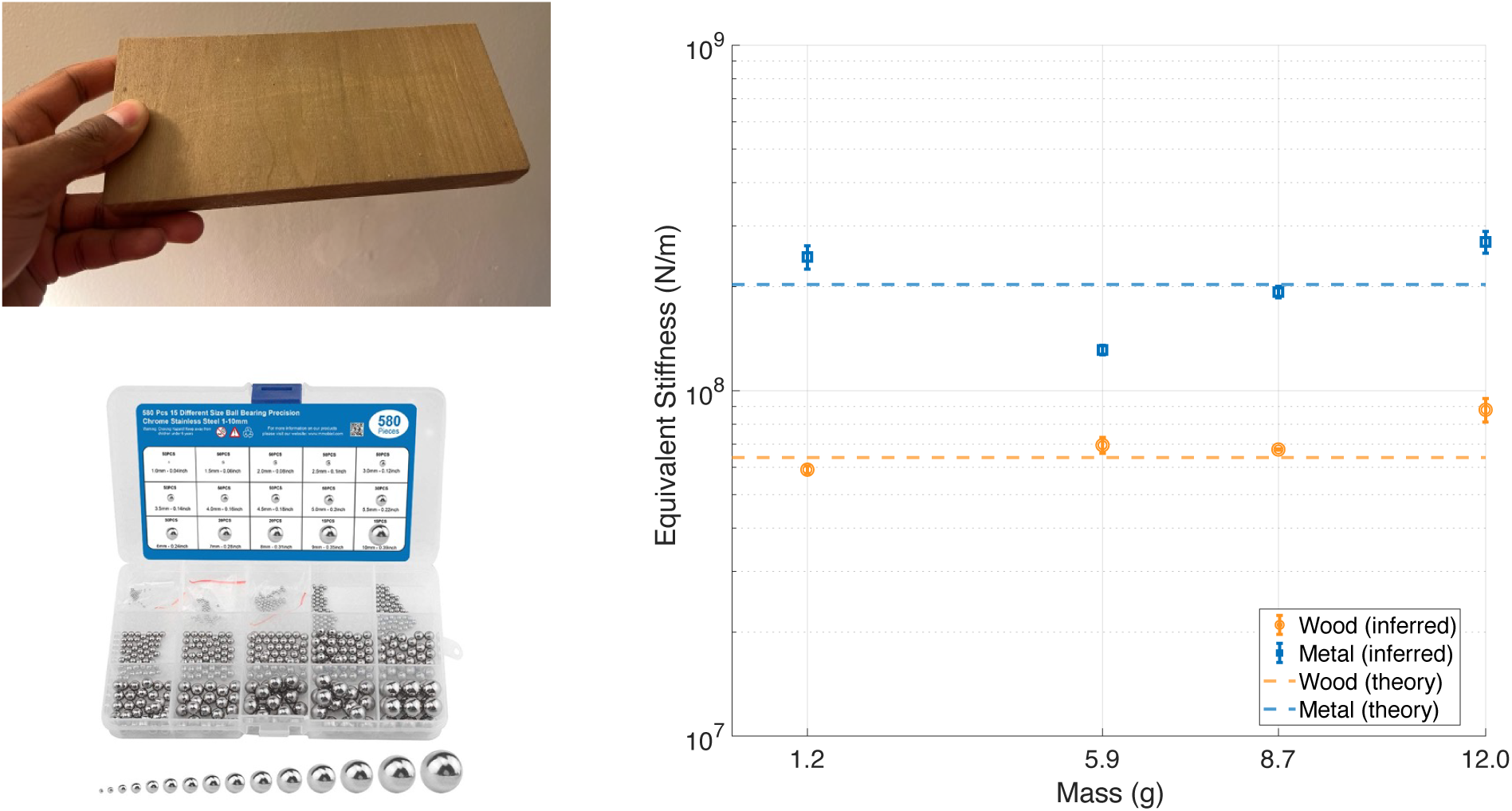
Validation of stiffness values and forcing function assumptions. Dots plot inferred stiffness for different impact masses and surface materials. Stiffness was inferred by fitting the generative model to the audio waveform recorded from impacts of small metal pellets on wood and metal surfaces. Dashed lines plot stiffness values used in the generative model for wood and metal in the small size category (given in Table 3 based on theoretical derivation using the constants in Table 2). See Methods for details.

